# Immunomodulatory RIG-I Agonist SLR14 Confers Protection Against a Range of Highly Pathogenic Negative Sense RNA viruses

**DOI:** 10.64898/2026.10.08.757743

**Authors:** Kristina Meier, Judith Olejnik, Joseline M. Velasquez-Reyes, Jonna B. Westover, Pushpinder Bawa, Elizabeth R. Wrobel, Corey W. Henderson, Pan Hu, Adam J. Hume, Alexander M. Holtz, Olga Fedorova, Gustavo Mostoslavsky, Darrell N. Kotton, John H. Connor, Andrea Marzi, Brian B. Gowen, Anna Marie Pyle, Elke Mühlberger

## Abstract

Emerging and re-emerging zoonotic RNA viruses, including filoviruses, henipaviruses and Rift Valley Fever virus (RVFV), pose a significant threat to human health. Due to their epidemic potential and high pathogenicity, these viruses are categorized as WHO priority pathogens, yet we still lack effective medical countermeasures (MCMs) against most of these dangerous pathogens, supporting that host-directed antivirals will be advantageous for broadly acting MCMs. Here we assessed the antiviral efficacy of SLR14, an immunomodulatory stem-loop RNA that activates RIG-I, for its activity against a panel of highly pathogenic viruses. SLR14 treatment provided broad-spectrum protection, markedly reducing infection with Ebola virus (EBOV), Marburg virus, and Sudan virus in human primary monocyte-derived macrophages, and Nipah virus replication in iPSC-derived endothelial cells, whether administered prophylactically or therapeutically. *In vivo*, a single dose of SLR14 protected mice against lethal RVFV infection when administered prophylactically, and, notably, retained significant efficacy when administered up to two days post-infection. Therapeutic SLR14 administration also improved survival rates of EBOV-infected mice. Although EBOV infection and SLR14 treatment activated overlapping innate immune pathways in human macrophages, viral infection induced a nonprotective response, whereas SLR14 elicited a well-regulated, markedly stronger and protective interferon-stimulated and inflammatory gene signature while avoiding prolonged IFN gene upregulation. Our data identify SLR14 as a potent host-directed therapeutic with broad antiviral efficacy against some of the most lethal viral pathogens.

## Introduction

Host-directed therapies (HDTs) against virus infection provide an advantage for pandemic preparedness because they avoid the need for pathogen identification and pathogen-specific treatment designs. Such pathogen-agnostic approaches are desirable in cases where infections are detected with a previously unknown pathogen. Since HDTs are designed to target universal responses, the innate immune system, our first line of defense against incoming pathogens, is an attractive target. Among other stimuli, innate immune responses are triggered by pattern recognition receptors (PRRs) that detect pathogen-associated molecular patterns (PAMPs). Retinoic acid-inducible gene-I (RIG-I) is a key sensor of viral infection that recognizes viral RNA and is ubiquitously expressed in almost all cell types. Binding of RIG-I to blunt-ended double-stranded RNA (dsRNA) with 5’ di- or -triphosphates triggers a signaling cascade leading to the induction of pro-inflammatory cytokines, chemokines, and type I interferons (IFNs-I), such as IFNα and IFNβ. Once secreted, IFNs-I act in an autocrine and paracrine manner to induce the upregulation of a plethora of interferon-stimulated genes (ISGs) to establish a potent antiviral state in infected and non-infected bystander cells (*1, 2*). RIG-I activation has emerged as a promising therapeutic strategy across multiple applications including cancer immunotherapy, vaccine adjuvancy, and broad-spectrum antiviral therapy (reviewed in (*3–9*)). RIG-I agonists with therapeutic potential include synthetic triphosphate dsRNAs such as 3pRNA (*10–16*), virus-derived RNAs such as hepatitis C-derived RAR/XRAR (*17–19*), SARS-CoV-2-derived nCov-L, VSV-derived 5’pppRNA (*9*), or Sendai virus defective interfering RNA (*20*).

A promising RIG-I agonist is stem-loop RNA 14 (SLR14), a small 5’-terminally diphosphorylated RNA that forms a stable blunt-ended hairpin structure with 14 base pairs, serving as a minimal RIG-I ligand (*21, 22*). SLR14 has similar potency to dsRNAs flanked by a 5’-terminal triphosphate, as RIG-I selectively recognizes the β-phosphate that is present in both types of agonists (*21, 23, 24*). SLR14 binding to RIG-I leads to the activation of the transcription factors IRF3 and NFκB followed by a strong IFN-I response, triggering antiviral and antitumor activities in both cultured cells and animals (*22, 25–27*). A single dose of SLR14 was previously shown to protect mice against lethal SARS-CoV-2 infection (*28*). Given that a delayed and dysregulated IFN response is a general hallmark of many severe viral infections (*29, 30*), timely and well-regulated activation of RIG-I using immunomodulatory compounds such as SLR14 and other RIG-I agonists represents a promising broad-spectrum antiviral strategy (*12, 31, 32*).

Emerging and re-emerging zoonotic RNA viruses pose a serious threat to public health. A prime example is Ebola virus (EBOV), the most prominent member of the *Filoviridae* family within the order *Mononegavirales*. Filoviruses are enveloped filamentous viruses that possess a non-segmented, negative-sense RNA genome and cause frequent outbreaks of severe disease with case fatality rates ranging from 40% to 90%. EBOV disease (EVD) is characterized by severe systemic infection with high viral titers, immune dysregulation, and coagulation abnormalities, ultimately leading to organ failure that is seen in late stages of fatal cases (*33*). Although an FDA-approved vaccine and monoclonal antibody treatments are available for EBOV (*34–36*), these medical countermeasures are highly specific to EBOV but not effective against other human-pathogenic filoviruses, such as Sudan virus (SUDV), Bundibugyo virus, or the more distantly related Marburg virus (MARV), all of which cause severe disease with high fatality rates similar to EVD. The innate immune system, specifically the IFN-I response, plays a decisive role in the clinical outcomes of filovirus disease. Although the pathogenic filoviruses efficiently counteract IFN-I induction and signaling in infected cells (*37*), filovirus infection still induces an IFN-I and inflammatory response in patients. However, these responses are not protective (*38, 39*).

Interference with innate immune signaling is also observed in infections with other highly pathogenic zoonotic viruses, such as Nipah virus (NiV; genus: *Henipavirus*, family: *Paramyxoviridae, order: Mononegavirales*), which causes almost annual outbreaks of severe respiratory disease and encephalitis in humans in Southeast Asia (*40, 41*). Like filoviruses, NiV belongs to the group of nonsegmented negative sense RNA viruses and is of zoonotic origin, with distinct fruit bats serving as its natural reservoir hosts (*40, 42*). Like filoviruses, NiV infection induces a nonprotective innate immune response (*43*), and it has been discussed that viral IFN-I antagonists may play a crucial role in immune dysregulation (*41, 44*). Due to the high case fatality rates caused by infection with these viruses along with their epidemic potential and severeness of disease, EBOV, MARV, SUDV, and NiV are classified as risk group 4 pathogens and listed as priority pathogens for research and development by WHO (*45*).

Rift Valley fever virus (RVFV; genus: *Phlebovirus*, family: *Phenuiviridae,* order: *Hareavirales*) is a mosquito-borne phlebovirus with a tripartite negative sense RNA genome that severely affects livestock, such as sheep and cattle. RVFV is endemic throughout Africa and the Arabian Peninsula and can be transmitted to humans with a variety of clinical outcomes, ranging from a self-limiting febrile illness to life-threatening disease (*46, 47*). Like the filoviruses and NiV, RVFV has been identified by WHO as a re-emerging pathogen of major public health concern (*45, 47*).

A common theme for emerging and re-emerging highly pathogenic viruses with epidemic potential is the lack of effective FDA-approved medical countermeasures. Developing novel, broad-acting and rapidly deployable antiviral strategies is essential for counteracting a wide variety of pathogenic viruses. In this study, we assessed the antiviral efficacy of SLR14 against various highly pathogenic negative sense RNA viruses. We show that SLR14 restricts the replication of filoviruses in human primary monocyte-derived macrophages (MDMs) by inducing a potent IFN-I-based innate immune response. Additionally, SLR14 treatment protected induced pluripotent stem cell (iPSC)-derived endothelial cells (iECs) against NiV infection. We further report that SLR14 pre-treatment protected mice against lethal RVFV infection and improved RVFV disease survival rates when administered up to 48 hours post-infection (hpi). Treatment with SLR14 at 8 hpi also enhanced survival rates in EBOV-infected mice. Taken together, our data show that SLR14 is an effective host-targeted antiviral with broad therapeutic efficacy against distinct segmented and nonsegmented negative-sense RNA viruses. Importantly, the broad antiviral spectrum of SLR14 demonstrated in this study suggests that SLR14 may also confer protection against other viruses from different viral families and orders.

## Results

### SLR14 induces a pronounced IFN-I response in human monocyte-derived macrophages

Macrophages are highly susceptible to filovirus infection and contribute to filovirus pathogenesis (*48*). Human MDMs are activated when infected with EBOV as reflected by early IRF3 nuclear translocation and a pronounced IFN-I and proinflammatory response (*49–51*). Thus, MDMs are an important *in vitro* model for studying filovirus infection in human primary cells. To define the innate immune response induced by SLR14 in these cells, we transfected MDMs from three donors with SLR14-diphosphate (SLR14; biologically active) to stimulate RIG-I. As controls, we treated the cells with transfection reagent only (TRO) or used SLR14 versions with a lower binding affinity for RIG-I that do not induce downstream signaling: SLR14-monophosphate (SLR14-mP; biologically inactive) and SLR14-OH (biologically minimally active) (*24*). Total cellular RNA was harvested at various time points post-transfection and used for bulk RNA-Seq analysis. Principal component analysis (PCA) showed that while SLR14 treatment induced significant differential gene expression (DGE) compared to untreated cells, the TRO control and the cells transfected with SLR14-OH or SLR14-mP clustered with untreated samples (**Fig. 1A**). Gene Set Enrichment Analysis (GSEA) on Hallmark gene sets revealed a strong induction of IFN and inflammatory response pathways upon SLR14 treatment at all analyzed time points with the highest upregulation of genes in these pathways detected at 9 hours post-transfection (**Fig. 1B**). At this time point, the top 10 upregulated genes included 5 different IFNs with the strongest upregulation seen for IFNβ1 (15.35 log2FC), followed by IFNL1 (14.53 log2FC). Overall, 18 IFNs were upregulated, peaking at 9 hours, with the highest expression levels observed for IFN β1, IFNL, IFNA1, IFNA2, IFNA8, IFNA13, IFNA14 and IFNW1 (**Fig. 1C and D**). In line with IFN induction, upregulation of multiple interferon-stimulated genes (ISGs) was observed (**Fig. 1C and E**). Expression of most ISGs increased over the observed time course with highest levels detected at 9 and 24 hours post-transfection. Upregulated ISGs included RNA sensors that activate inflammatory or cell death signaling (IFIT1, ZBP1, AIM2, IFIT2), inhibitors of viral replication (RSAD2/Viperin), proteins leading to viral RNA degradation or modification (ISG20, APOBEC3A), and inhibitors of viral entry (IFIT1M, CH25H) (**Fig. 1E**). Aconitate decarboxylase 1 (ACOD1/IRG1), a regulator of inflammation and immunometabolism that is upregulated in a variety of bacterial and viral diseases (*52*), was most highly expressed by 3 hours after SLR14 treatment (**Fig. 1E**). We also observed strong chemokine upregulation, specifically for the IFN-inducible CXCL9 family (CXCL9, 10 and 11), whose members were among the top 10 DEGs (13.41 log2FC for CXCL9, 18.53 log2FC for CXCL10 and 16.72 log2FC for CXCL11 at 9 hours), and multiple members of the CC chemokine ligands with the highest upregulation observed for CCL8 (**Fig. 1C and F**, CCL8, CCL2L1, CCL4, CCL4L2). In addition, we detected cytokine upregulation, including members of the IL12 family (IL12A, IL12B, and IL27) and TNF signaling (TNF, TNFSF10, TNFSF18). However, the induction of these genes was generally lower compared to the upregulated chemokines. Cytokine expression peaked at 9 hours, as observed for CXCL9 and CXCL10, whereas CXCL11 and CCL8 reached the highest levels at 24 hours (**Fig. 1F**).

**Figure 1.**
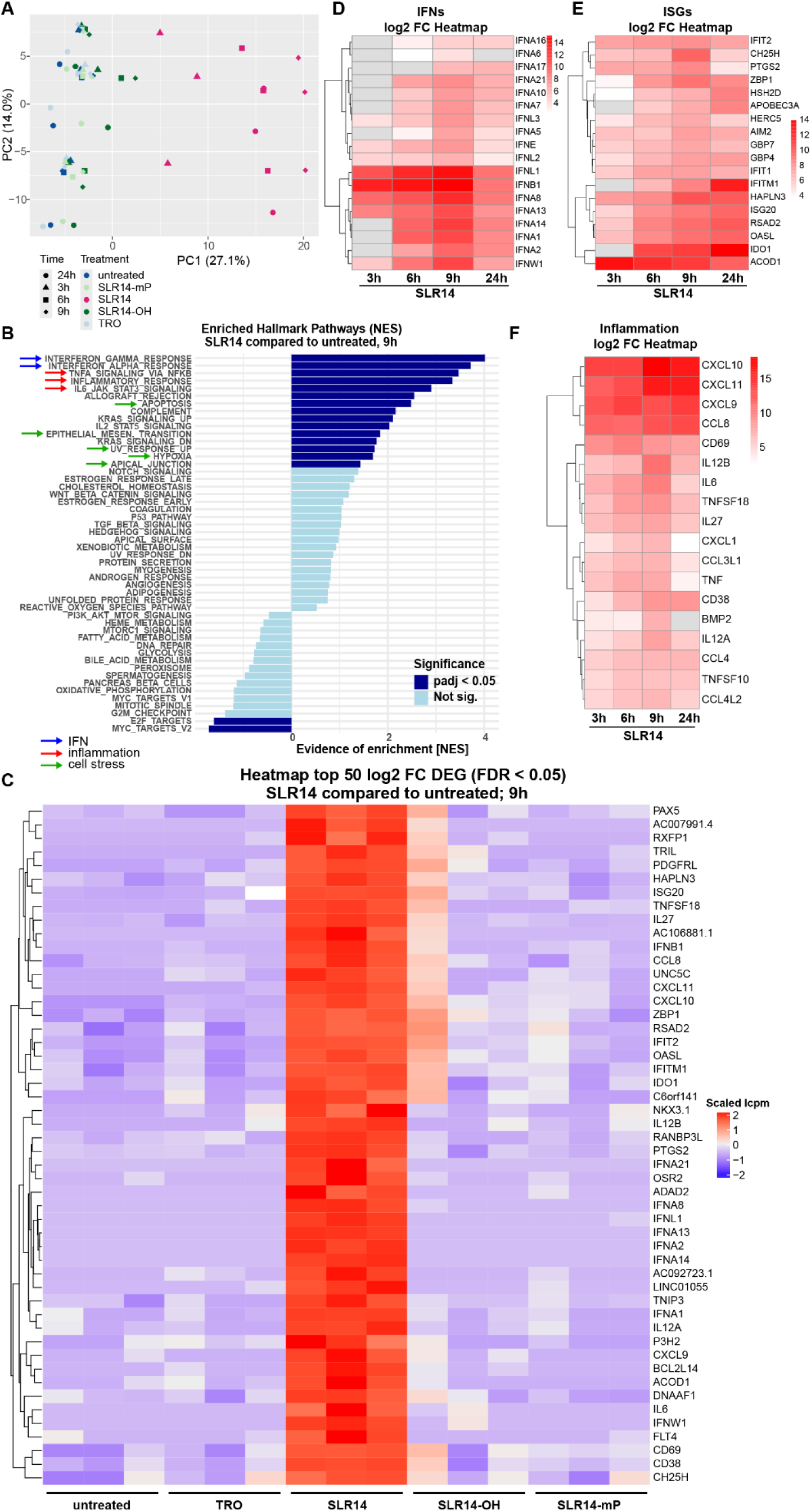
SLR14-induced IFN-I response in human monocyte-derived macrophages. MDMs from three different donors seeded in 24-wells were transfected with 1 μg SLR14, 1 μg SLR14-mP, 1 μg SLR14-OH, transfection reagent only (TRO), or left untreated. Total cellular RNA was harvested at the indicated time points for RNA-Seq analysis. (**A**) Principal component analysis (PCA) of the transcriptomic responses in MDMs to the respective treatments. Different treatment conditions marked by colors, different time points marked by symbols. (**B**) Gene set enrichment analysis (GSEA) using the Hallmark gene sets on the top differentially expressed gene (DEG) sets in SLR14-treated MDMs at 9 hours post-transfection compared to untreated controls. Light blue bars, not significant; dark blue bars, statistically significant (p < 0.05). (**C**) Heatmap of top 50 log2 FC genes of SLR14-treated MDMs compared to untreated cells. Replicates represent the three donors. (**D**) Log2 FC heatmap of IFN genes differentially expressed at different time points after SLR14 treatment compared to untreated cells. (**E**) Log2 FC heatmap of ISGs in top 100 DEGs at 9 hours of SLR14-treated samples compared to untreated cells across time points. (**F**) Log2 FC heatmap of inflammatory mediators in top 110 DEGs at 9 hours of SLR14-treated samples compared to untreated cells across time points. The data from three donors are combined.

In contrast to SLR14, the SLR14-OH and SLR14-mP controls did not lead to significant activation of the pathways induced by SLR14 treatment at any time point analyzed (**Fig. 1C and S1**). Low induction of IFN genes and ISGs in the MDMs from one donor was observed in SLR14-OH treated samples, which might be due to residual but inefficient RIG-I activation (*24*). The effect of SLR14-mP was even more subtle, consistent with previous observations indicating that 5’-monophosphorylated RNAs may even block RIG-I activation (**Fig. 1C and S1**) (*24*). Overall, our data demonstrate that SLR14 is a potent activator of RIG-I in MDMs with subsequent strong upregulation of both IFN and inflammatory signaling cascades.

### EBOV infection of MDMs induces ISG expression mainly in uninfected bystander cells

Although EBOV is sensitive to IFN-I treatment (*53–55*), the IFN response observed in EBOV-infected patients is not protective (*38, 56*). This might be due to the ability of EBOV to suppress IFN-I induction and signaling in infected cells, leading to delayed and/or dysregulated innate immune responses (reviewed in (*37, 57*)).

While many cell types infected with EBOV display a negligible or moderate delayed IFN-I response (*58–61*), macrophages including human MDMs are strongly activated when infected with EBOV (*49–51, 62–64*). To be able to compare the innate immune responses induced by EBOV to those triggered by SLR14 treatment in our MDM model, we performed bulk RNA-Seq on EBOV-infected MDMs. EBOV infection was the primary driver of variance in the dataset, clearly separating infected samples from mock controls in PCA. Secondary differences among infected samples were attributable to donor variation (**Fig. 2A**). GSEA showed a significant upregulation of genes associated with IFN response pathways, inflammation, and cell stress in EBOV-infected cells (**Fig. 2B**). Similar to SLR14-treated MDMs, ISGs (e.g., RSAD2/Viperin, HSH2D, ZBP1, SerpinB2, IDO1, ISG20), IFN genes (e.g., IFNB1, IFNL1) and chemokines (e.g., CXCL8, CXCL11) dominated the top 50 differentially expressed genes in EBOV-infected MDMs across all three donors compared to non-infected controls (**Fig. 2C and D**). Overall, more transcripts were upregulated than downregulated during EBOV infection, with viral genes among the most highly expressed (**Fig. 2D**).

**Figure 2.**
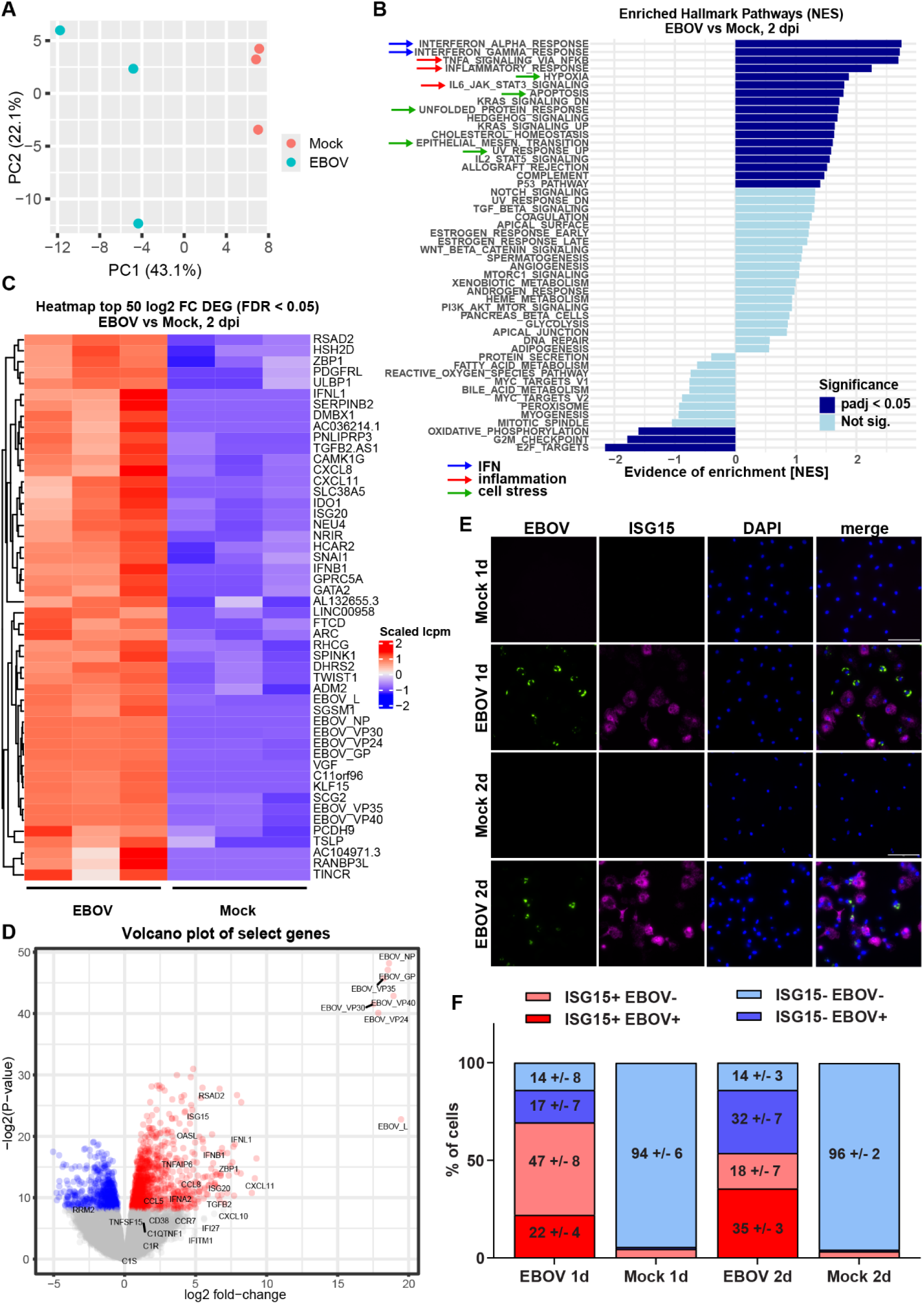
Innate immune responses induced by EBOV infection of MDMs. **(A-D)** MDMs from three different donors seeded in 24-wells were infected with EBOV at MOI 1. 2 dpi, total cellular RNA was harvested and used for bulk RNA-Seq. (**A**) PCA of the transcriptomic response to EBOV infection of human MDMs. (**B**) GSEA using the Hallmark gene sets on the top DEG sets in EBOV-infected MDMs compared to mock-infected controls. Light blue bars, not significant; dark blue bars, statistically significant (p < 0.05). (**C**) Heatmap of top 50 log2 FC genes of EBOV-infected MDMs compared to mock-infected controls. Replicates represent the three donors. (**D**) Volcano plot of DEGs in EBOV-infected MDMs compared to mock-infected controls. Viral genes and select IFNs and ISGs are labeled. (**E**) MDMs from three different donors seeded in 8-well chamber slides were infected with EBOV at MOI 10 and fixed at 1 or 2 dpi. RNA FISH analysis was performed with probes directed against EBOV genomic RNA (green) and ISG15 mRNA (magenta). Nuclei were stained with DAPI (blue). Scale bars, 100 µm. Representative images are shown. (**F**) Quantification of EBOV- and ISG15-positive cells from multiple images (n ≥ 6 per donor) of MDMs from three donors using QuPath. Percentages of cell populations + SD are shown.

Since bulk RNA-Seq analysis does not capture the spatial transcriptomic profile of infected and non-infected bystander cells, we performed RNA fluorescence in situ hybridization (FISH) to determine ISG expression at single cell level. MDMs were infected with EBOV, fixed at 1 or 2 days post-infection (dpi), and analyzed by RNA FISH using probes binding to the EBOV genome and ISG15 mRNA (**Fig. 2E**). Only a small fraction of mock-infected cells expressed ISG15 at 1 or 2 dpi, and these cells formed isolated clusters in some of the areas imaged. In the EBOV infections, 69% of the cells were positive for ISG15 mRNA and 39% for EBOV RNA at 1 dpi (**Fig. 2E and F**). 68% of ISG15^+^ (47% of total cells) were negative for EBOV RNA, indicating that ISG expression was induced mainly in non-infected bystander cells. At 2 dpi, the number of EBOV^+^ cells increased to 67%, with 35% EBOV^+^/ISG15^+^ cells and 32% of EBOV^+^/ISG15^-^ cells (**Fig. 2F**). The percentage of EBOV^-^/ISG15^+^ decreased (18%) compared to 1 dpi (47%). These results suggest that while non-infected bystander cells were the main mediators of the observed IFN response in MDMs, particularly at 1 dpi, the induced ISG response was not sufficient to restrict EBOV replication as infection rates continued to increase at 2 dpi.

### Prophylactic treatment of MDMs with SLR14 inhibits EBOV and SUDV infection

Since SLR14 induces a strong, well-controlled early RIG-I-mediated innate immune response in human MDMs (**Fig. 1**), we next explored if prophylactic treatment with SLR14 would be protective against infection with EBOV, and a second distinct filovirus, SUDV. MDMs were transfected with SLR14, SLR14-OH, treated with TRO, or left untreated. At 6 hours post-transfection, the cells were infected with EBOV or SUDV at an MOI of 10 or 1, fixed 1 or 2 dpi, and infection rates were determined by immunofluorescence analysis (IFA). We observed a clear reduction of EBOV- and SUDV-positive cells in the SLR14-treated samples compared to the controls (**Fig. 3A and B**). At 1 dpi, approximately 10% of the SLR14-treated MDMs were infected with EBOV or SUDV, compared to approximately 45% of untreated cells or the TRO controls, indicating that prophylactic SLR4 treatment potently protects MDMs from EBOV and SUDV infection (**Fig. 3C and D**). The protective effect was even more pronounced at 2 dpi in cells infected with an MOI of 1, with an average infection rate of 19% in SLR14-treated cells compared to 80% in control cells in the case of EBOV, and approximately 5% in SLR14-treated cells infected with SUDV compared to 79% or 88% in untreated cells or cells treated with TRO, respectively (**Fig. 3**). The control RNA SLR14-OH, which induced no or minor activation of IFN signaling pathways (**Fig. 1 and S1**), still conferred a limited level of protection, albeit less than the biologically active SLR14. Similar results were obtained using recombinant EBOV expressing ZsGreen as a reporter gene (*65*), with the percentage of fluorescent cells serving as the readout for infection rates (**Fig. S2**).

**Figure 3.**
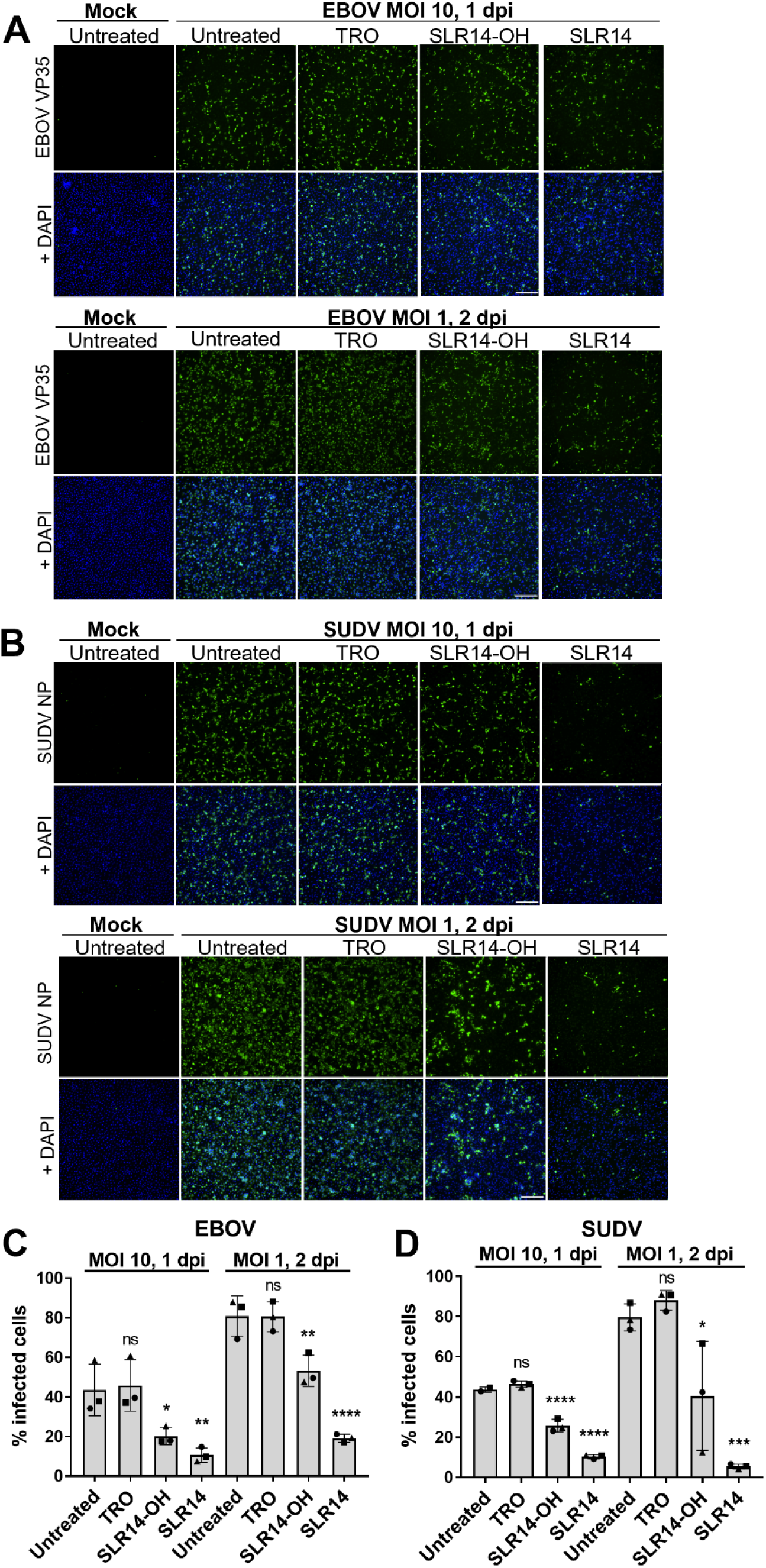
Prophylactic treatment of MDMs with SLR14 inhibits EBOV and SUDV infection. **(A, B)** MDMs from three different donors seeded in 24-wells were transfected with 1 μg SLR14, 1 μg SLR14-OH, transfection reagent only (TRO), or left untreated. 6 hours post-transfection, the cells were mock-infected or infected with EBOV **(A)** or SUDV **(B)** at MOI 10 or 1 and fixed at 1 dpi or 2 dpi. The cells were stained with antibodies against EBOV VP35 (green) or SUDV nucleoprotein (NP, green). Cell nuclei were stained with DAPI (blue). Representative immunofluorescence images are shown. Scale bars, 250 µm. **(C, D)** Infection rates of EBOV **(C)** or SUDV **(D)** as quantified from multiple images (n ≥ 6 per donor) of MDMs from three donors using QuPath. Donors represented by different symbols. Significant differences between conditions were determined by one-way ANOVA followed by Dunnett’s post-test compared to Untreated control. P-values: *<0.1; **<0.01; ***<0.001; ****<0.0001; ns: not significant.

Next, we assessed whether there are differences that define a protective SLR14-induced immune response compared to a non-protective virus-induced response. To this end, MDMs were transfected with TRO, SLR14-OH or SLR14, or left untreated. At 6 hours post-transfection, the cells were infected with EBOV or SUDV at MOI 1, and total RNA was harvested at 2 dpi for bulk RNA-Seq analysis. PCA showed that SLR14 treatment overwhelmingly structured the transcriptional landscape, producing the primary axis of separation, with viral identity (EBOV versus SUDV) imposing a strong secondary clustering effect. (**Fig. 4A**). A similar trend was seen in MDMs transfected with SLR14 and subsequently infected with SUDV at an MOI of 10 and harvested at 1 dpi with SLR14-treated samples clustering separately from untreated infected samples (**Fig. S3A**). GSEA revealed that SLR14 treatment elicited a transcriptomic response in EBOV-infected cells that closely mirrored the response observed in SLR14-treated non-infected cells (**Fig. 4B** and **1B**), with strong upregulation of ISGs and chemokines/cytokines in SLR14-treated EBOV-infected MDMs (e.g., IFITM1, RSAD2, OASL, ISG20, ZBP1, ACOD1, CD38,

**Figure 4.**
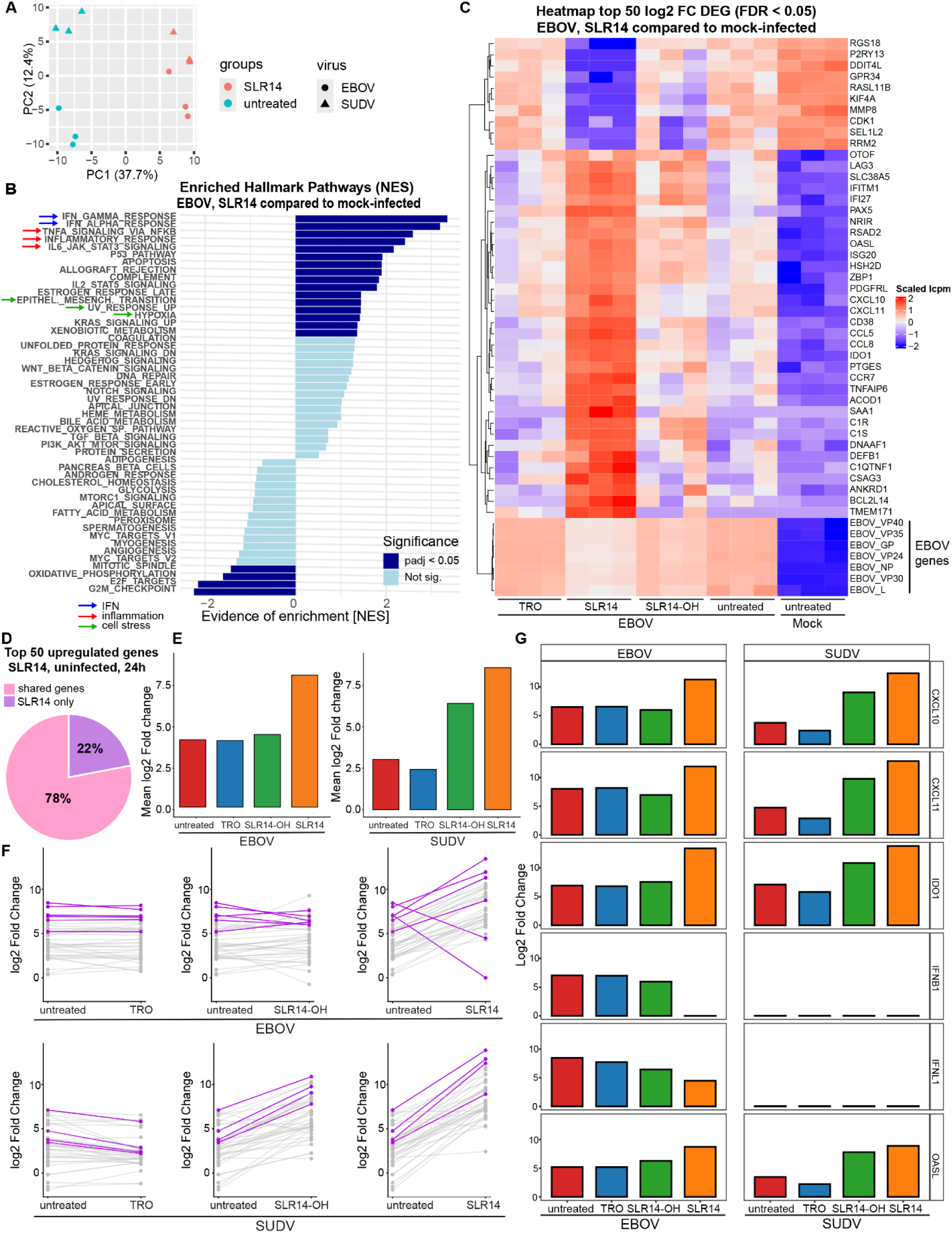
SLR14 treatment induces IFN and inflammatory responses in filovirus-infected MDMs. MDMs from three different donors seeded in 24-wells were transfected with 1 μg SLR14, 1 μg SLR14-OH, transfection reagent only (TRO), or left untreated. 6 hours post-transfection, the cells were infected with EBOV or SUDV at MOI 1 and harvested at 2 dpi. Total cellular RNA was isolated and used for bulk RNA-Seq analysis. (**A**) PCA of the transcriptomic response of the to the respective treatments. EBOV and SUDV infection marked by symbols, different treatment conditions marked by colors. (**B**) GSEA using the Hallmark gene sets on the top DEG sets in SLR14-treated, EBOV-infected MDMs compared to untreated, mock-infected controls. Light blue bars, not significant; dark blue bars, statistically significant (p < 0.05). (**C**) Heatmap of top 50 log2 FC genes in SLR14-treated, EBOV-infected MDMs compared to mock-infected controls for all tested treatment conditions. Replicates represent the three donors. (**D**) Pie chart visualizing the genes shared between the datasets (pink) that are upregulated in SLR14 only experiments and those specific to SLR14 only (purple). (**E**) Average log2 FC value of the 39 shared genes in the datasets. (**F**) Slope plot of log2 FC of the 39 shared genes from (D) in untreated EBOV- or SUDV-infected samples compared to TRO-, SLR14-OH-, and SLR14-treated samples. The genes with the largest expression difference between untreated and SLR14-treated infected cells are highlighted in magenta in each plot. (**G**) Comparison of genes with the largest expression difference between SLR14 treatment and all other conditions.

CXCL10, CXCL11, CCL8, **Fig. 4C**). This was also true for SUDV-infected cells pretreated with SLR14, irrespective of MOI used or time point (**Fig. S3B and S3C**). Upregulation of select genes was verified by RT-qPCR (**Fig. S4**). Downregulated genes in SLR14-treated EBOV-infected (**Fig. 4B and C**) or SUDV-infected MDMs (**Fig. S3**) also reflected observations of SLR14-treated non-infected cells (**Fig. 1B and S1**). Importantly, RNA-Seq analysis validated the antiviral effect of SLR14, revealing reduced levels of viral transcripts in MDMs pre-treated with SLR14 for 6 hours prior to infection with EBOV or SUDV (**Fig. 4C and S3**). EBOV infection alone elicited an overall transcriptional response in MDMs similar to that observed with SLR14 treatment in non-infected cells (compare **Fig. 2B and C** to **Fig. 1B and C**).

To identify potential differences between the protective host response elicited by SLR14 treatment and the non-protective response induced by viral infection, we performed a second analysis focusing on genes that exhibited the largest changes in expression (log2 fold change > 7) in uninfected, SLR14-treated MDMs at 24 hours (**Fig. S5A**). The expression levels of these 50 genes were compared to EBOV- or SUDV-infected samples across all treatment conditions (untreated, TRO, SLR14-OH, SLR14). Of the 50 most highly upregulated genes in uninfected, SLR14-treated MDMs (**Fig. S5A**), 39 (78%) were also upregulated in the EBOV- and SUDV-infected cells regardless of treatment conditions (**Fig. 4D**). Interestingly, the majority of the 11 genes (22%) only upregulated in the uninfected, SRL14-treated cells belong to the group of IFNA and IFNW genes (**Fig. 4D and S5B**). This difference in expression levels could be due to different sampling times (24 hours versus 48 hours). Among the shared genes, numerous ISGs and cytokine- and chemokine-related genes were upregulated across all conditions following viral infection (**Fig. S5C and D**). Averaging the expression levels of the 39 shared genes revealed that the magnitude of gene induction was the lowest in untreated EBOV- and SUDV-infected cells and the TRO controls (**Fig. 4E**). While the infected SLR14-OH-treated samples displayed an intermediate phenotype, specifically in the SUDV infections, the most pronounced changes in expression levels were consistently found in the infected SLR14-treated samples (**Fig. 4E)**. To better understand gene expression dynamics, we compared the log2 fold changes (LFC) of the 39 individual genes in TRO-, SLR14-OH- or SLR14-treated infected samples to untreated infected samples. While the LFC of the individual genes in EBOV-infected cells treated with TRO or SLR14-OH were similar to those in untreated cells, all but two genes were more highly upregulated in EBOV-infected SLR14-treated cells compared to untreated cells (**Fig. 4F).** A similar trend was observed in SUDV-infected cells, with the transcriptional response of SLR14-OH-treated cells falling between those of the other conditions (**Fig. 4F**). The genes exhibiting the greatest absolute differences between conditions are highlighted in purple. Among the highest upregulated genes in SLR14-treated EBOV- or SUDV-infected samples were the cytokines CXCL10, CXCL11, as well as the macrophage activation marker IDO1. All three reached LFC of 10 or higher, whereas induction in the untreated and control groups was considerably lower (LFC of 8 to 6) (**Fig. 4G**). A similar trend was observed for the classic IFN responsive gene OASL showing the highest upregulation in SLR14-treated infected cells compared to all other conditions (**Fig. 4G**). Strikingly, we did not observe a significant upregulation of the IFN genes IFNB1 and IFNL1 in SLR14-treated cells infected with EBOV at 2 dpi, whereas these genes were highly upregulated in untreated EBOV-infected MDMs and the control groups (**Fig. 4G**). Interestingly, there was no upregulation of IFNB1 or IFNL1 in any samples infected with SUDV at 2 dpi (**Fig. 4G)**. These results likely reflect divergent IFN expression dynamics between different filoviruses, suggesting that the peak upregulation of IFN genes in SUDV infection may occur outside the single 2 dpi time point we analyzed here.

Collectively, our findings show that although both filoviral infection and SLR14 treatment trigger the induction of similar innate immune signatures, the SLR14-induced response is defined by a stronger and earlier upregulation of ISGs and cytokines compared to viral infection alone. These results indicate that a potent anti-filoviral response depends on both robust amplitude and coordinated activation of innate immune mediators, while limiting prolonged upregulation of IFN genes to prevent excessive inflammation and immune dysregulation.

### SLR14 has broad antiviral efficacy against distantly related filoviruses when administered post-infection

We next assessed the protective effect of SLR14 against filovirus infection when administered post-infection. In addition to EBOV and SUDV, we included the more distantly related MARV in this experiment. Since we observed that SLR14-OH had an intermediate effect in the previous experiment, we used SLR14-mP as a control going forward. MDMs from at least three different donors were infected with EBOV, SUDV or MARV at an MOI of 0.01. At 1 dpi, the cells were transfected with SLR14, SLR14-mP, TRO, or left untreated. Cell culture supernatants and total cellular RNA were collected, and the remaining cells were fixed at day 7 post-infection. EBOV infection rates were determined by RT-qPCR quantification of viral transcripts, while SUDV and MARV infection rates were determined by immunofluorescence analysis. We found that SLR14 treatment significantly reduced viral spread compared to cells that were treated with the control SLR14-mP or TRO for all three tested viruses (**Fig. 5A-F**). We also observed that SLR14 treatment led to a significant reduction of viral titers (at least 2 logs less) in cell culture supernatants of MDMs infected with EBOV, SUDV or MARV (**Fig. 5G**). Similar results were obtained using recombinant EBOV-ZsGreen, with infection rates determined by quantifying ZsGreen-expressing cells (**Fig. S6A and B**). To determine the temporal window of SLR14 efficacy, we infected MDMs with EBOV-ZsGreen and treated the cells with SLR14 at 1-4 dpi. SLR14 treatment up to 3 dpi significantly reduced infection rates with the strongest effects observed when SLR14 was administered at 1 or 2 dpi (**Fig. S6C**). Taken together, these data suggest that post-infection treatment with SLR14 broadly inhibits viral replication across different filovirus genera.

**Figure 5.**
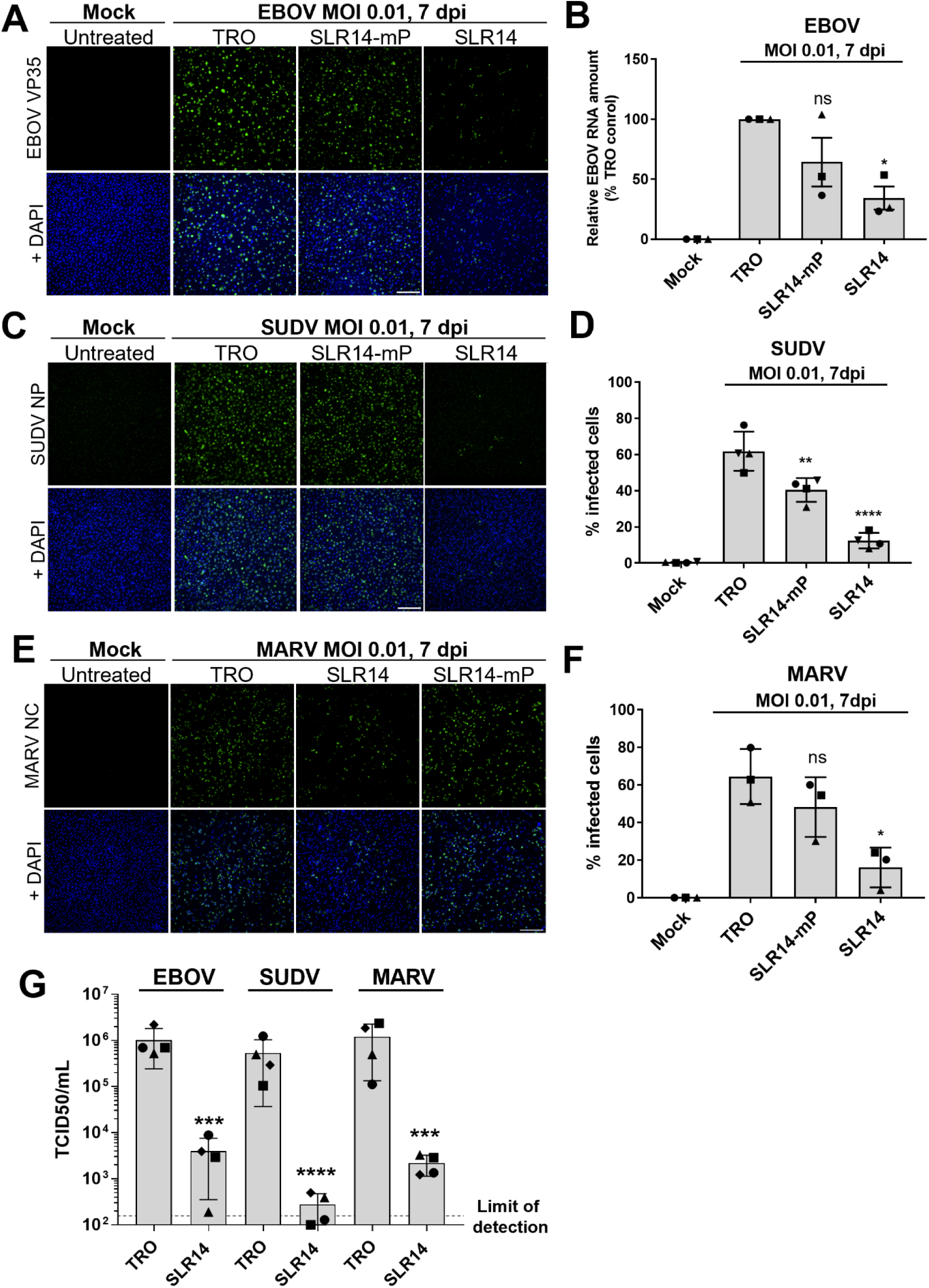
SLR14 restricts filovirus infection when administered post-infection. (**A, B**) MDMs from three or four donors seeded in 24-wells (3x10^5^/well) were mock-infected or infected with EBOV or (**C, D**) SUDV or (**E, F**) MARV at MOI 0.01. At 1 dpi, the cells were transfected with 1 μg SLR14, 1 μg SLR14-mP, transfection reagent only (TRO), or left untreated. At 7 dpi, total cellular RNA was isolated, or the cells were fixed and stained with antibodies against EBOV VP35 (**A**, green), SUDV NP (**C**, green), or MARV nucleocapsid (NC) (**E**, green). Cell nuclei were stained with DAPI (blue). Representative immunofluorescence images are shown. Scale bars, 250 µm. **(B)** Quantification of EBOV RNA by RT-qPCR using primers binding to the EBOV VP35 gene. Beta-2-microglobulin primers were used for normalization. VP35 gene expression shown as percent of TRO control. **(D)** SUDV and (**F**) MARV infection rates as quantified from multiple images (n ≥ 4 per donor) of MDMs from three (MARV) or four donors (SUDV) using QuPath. Donors represented by different symbols**. (G)** Viral titers determined by TCID_50_ assay from cell culture supernatants harvested before fixation of the MDMs at 7 dpi shown in (A-E). Viral titers of some SLR14-treated samples are below detection limit. Statistical analysis of conditions compared to TRO was performed by one-way ANOVA followed by Dunnett’s post-test (B, D, F) or unpaired two-tailed T-test (G). P-values: *<0.1; **<0.01; ***<0.001; ****<0.0001; ns: not significant.

### SLR14 inhibits Nipah virus infection

To assess the breadth of SLR14 antiviral effects, we tested the efficacy against the highly pathogenic NiV. Since endothelial cells are main target cells of NiV infection (*66*), we differentiated human induced pluripotent stem cells (iPSCs) into primary-like endothelial cells (iECs; F**ig. 6A**). iECs have previously been shown to be highly permissive to NiV infection (*67*) and were also permissive in our infection model (**Fig. 6B and S7A**). Previous work showed minimal changes in the transcriptomic profile of NiV-infected artery iECs at early time points of infection (5-15 hpi) (*67*). Similarly, we did not observe statistically significant changes in the transcriptomic profile of NiV-infected iECs at 6 hpi (**Fig. S7B and D**). However, at 24 hpi, we observed robust IFN and ISG expression in our iEC infection model, suggesting that NiV induces a delayed innate immune response in these cells (**Fig. S7B, C, E**). To assess ISG expression in NiV-infected iECs at single cell level, we performed RNA FISH analysis using probes targeting NiV L and ISG15 mRNA. Robust staining for NiV and ISG15 mRNA was observed in the NiV-infected samples, whereas ISG15 staining was largely absent in mock-infected cells except for infrequent, small clusters of ISG15-positive cells (**Fig. 6B**). Similar to our data obtained with EBOV-infected MDMs (**Fig. 2E**), ISG15 mRNA staining was observed mainly in uninfected bystander cells in the NiV-infected samples. Quantification of NiV L and ISG15 positive cells confirmed that most ISG-15-expressing cells were negative for NiV mRNA (74% of all ISG-positive cells) (**Fig. 6C**). Together, these data indicate that although NiV infection induces an innate immune response in iECs, this response is considerably delayed and non-protective.

**Figure 6.**
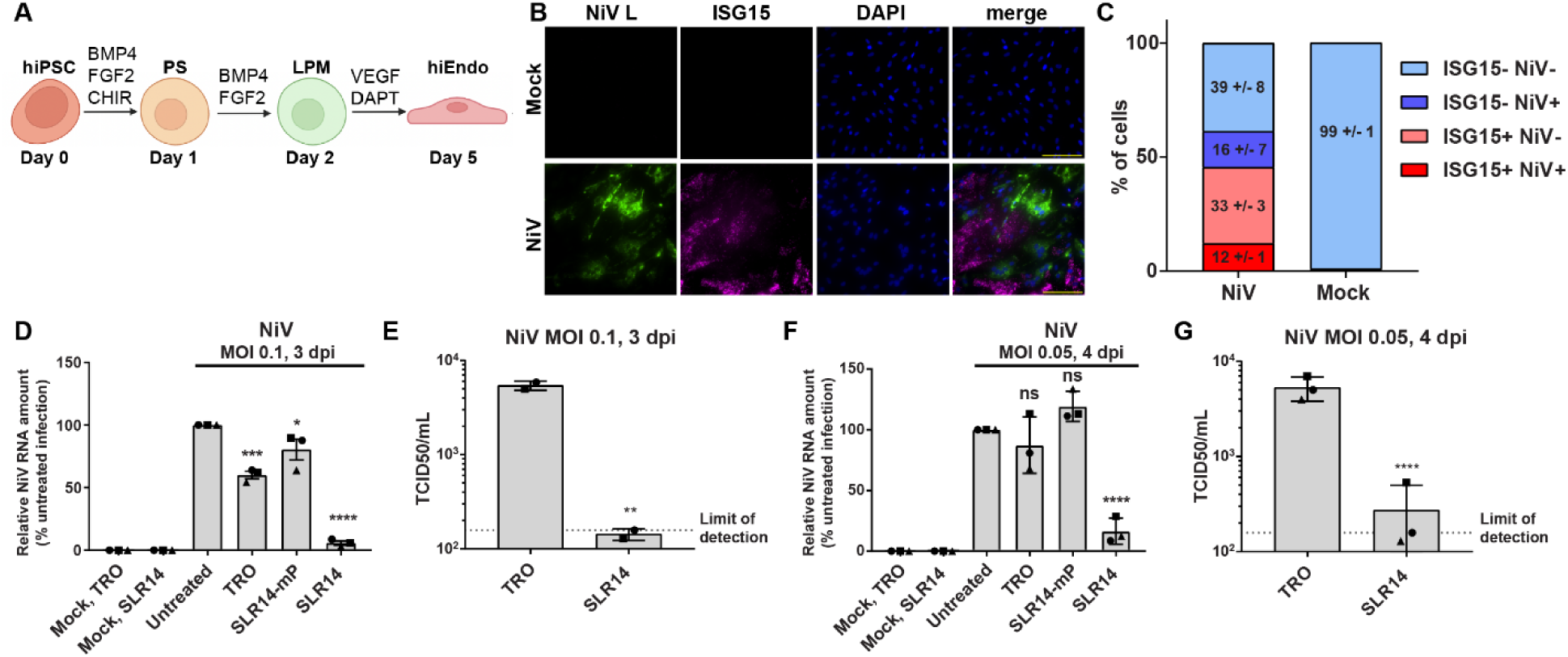
SLR14 protects human iECs against NiV infection. (**A**) Scheme of human iPSC differentiation into iECs. (**B**) RNA FISH analysis of NiV-infected iECs. iECs seeded in 8-well chamber slides were infected with NiV at MOI 10, fixed at 24 hpi, and stained with RNAscope probes directed against the NiV polymerase gene (L) to detect (+) sense RNA (viral mRNA and antigenomic RNA; green) and probes against ISG15 mRNA (magenta). Nuclei were stained with DAPI (blue). Scale bars, 100 μm, Representative images are shown. (**C**) Quantification of NIV- and ISG15-positive cells from multiple images (n ≥ 10 per experiment) of iECs from three independent experiments using QuPath. Percentage of cells populations +/SD are shown. (**D, E**) Prophylactic administration of SLR14 followed by NiV infection. iECs seeded in 24-wells (1.8x10^5^/well) were left untreated or transfected with 25 ng SLR14, 25 ng SLR14-mP, or transfection reagent only (TRO). 6 hours post-transfection, the cells were infected with NiV at MOI 0.1. Mock-infected cells were used as controls. (**D**) Total cellular RNA was harvested at 3 dpi and used for RT-qPCR with primers binding to the NiV nucleocapsid protein (N) gene. Beta-2-microglobulin primers were used for normalization. NiV N gene expression shown as percent of untreated control. Experiment performed in triplicate; symbols represent each experiment. (**E**) Cell supernatants were collected at 3 dpi and used for TCID_50_ assays to determine viral titers. Viral titers of some SLR14-treated samples are below detection limit. **(F, G**) Post-infection SLR14 treatment. iECs seeded in 24-wells (1.8x10^5^/well) were infected with NiV at MOI 0.05. At 1 dpi, the cells were left untreated or transfected with 25 ng SLR14, 25 ng SLR14-mP, or transfection reagent only (TRO). Mock-infected cells were used as controls. (**F**) Total cellular RNA was harvested 4 dpi and used for RT-qPCR as described in (D). Experiment performed in triplicate; symbols represent each experiment. (**G**) Cell supernatants were collected at 4 dpi and used for TCID_50_ assays to determine viral titers. Viral titers of some SLR14-treated samples are below detection limit. Statistical analysis of conditions compared to untreated control performed by one-way ANOVA followed by the Dunnett’s post-test (D, F) or unpaired two-tailed T-test (E, G). P-values: *<0.1; **<0.01; ***<0.001; ****<0.0001; ns: not significant.

We next performed SLR14 transfection experiments to assess whether SLR14 would induce a strong and early IFN response in iECs. We first evaluated iEC sensitivity to SLR14 treatment using a dose-response approach and found that even at a 40-fold lower amount (25 ng) than that used for MDMs, SLR14 still induced significant IFNB and TNFA mRNA expression (**Fig. S8A and B**).

IFNB was already induced at 3 hours post-transfection in SLR14-transfected iECs, with peak levels at 6 hours, whereas the controls showed no induction (**Fig. S8C**). Using 25 ng of SLR14, we tested its antiviral activity against NiV when administered either 6 hours pre- or 1 day post-infection. Briefly, iECs were transfected and 6 hours later infected with NiV at MOI 0.1. For post treatment studies, cells were infected at MOI 0.05 and transfected with 25 ng SLR14, 25 ng SLR14-mP or TRO, or left untreated at 1 dpi. At 3 dpi (pretreatment) or 4 dpi (post-infection treatment), cell culture supernatants and total RNA were harvested. Quantification of relative viral RNA amounts via RT-qPCR targeting the NiV nucleocapsid protein (N) gene demonstrated that SLR14 treatment of iECs either pre- or post-infection significantly reduced viral RNA levels compared to control cells (**Fig. 6D and F**). Additionally, we observed a reduction in viral titers in cell culture supernatants from SLR14-treated cells to almost detection limit or below **(Fig. 6E and G)**. This indicates that SLR14 is broadly active against different nonsegmented negative sense RNA viruses from distinct families. This observation is of particular interest since advances in high-throughput sequencing have led to the discovery of numerous novel filoviruses and henipaviruses with unknown pathogenic potential (*68, 69*).

### SLR14 protects mice against RVFV disease

To extend our *in vitro* findings to an *in vivo* model, we first assessed the protective efficacy of SLR14 against RVFV in a lethal murine infection model (*70*) under BSL-3 conditions. Initially, we tested whether SLR14 activated RIG-I when administered intraperitoneally (IP) in mice and determined high levels of circulating IFNα as measured 6 hours after SLR14 treatment (**Table S1**). To evaluate pre-exposure efficacy, doses of p3SLR14 (triphosphorylated SLR14, see Methods) or vehicle control were administered IP to BALB/c mice, and 6 hours later, the mice were infected with RVFV. Mice were monitored daily for weight loss and survival for up to 21 dpi. Complete protection was observed using 25 μg of p3SLR14, and, remarkably, 90% of the animals survived when treated with 1 or 5 µg of SLR14 (**Fig. 7A and B**). In contrast, 70% of the vehicle-treated mice succumbed to infection (**Fig. 7A and B**). In practice, antivirals are administered post-exposure or therapeutically when the patient presents with clinically apparent disease. Therefore, we evaluated post-infection intervention with SLR14 following RVFV challenge. BALB/c mice were infected with RVFV and subsequently treated IP with 5 µg SLR14 at 8, 24 or 48 hours after virus challenge. All PBS control animals and 7 out of 8 animals of the vehicle control group succumbed to infection, while SLR14 post-infection treatment protected 50% of mice against RVFV in all treatment groups with prolonged survival when SLR14 was administered at earlier time points (**Fig. 7C and D**). Four animals per treatment group were euthanized at 3 dpi to determine RVFV titers in serum and tissue. Early SLR14 treatment at 8 hpi significantly reduced viral load across serum, liver, and spleen (**Fig. 7E**). However, viral titers were not significantly reduced when treatment was administered at later time points, despite substantially improved survival rates. The lack of viral reduction at later time points likely reflects variable protective efficacy within treatment groups. Given the 50% survival rate, some animals might not have mounted adequate antiviral immunity at the time of sampling and subsequently would have succumbed to infection. This is consistent with the considerable variability in viral titers across animals within the treatment groups (**Fig. 7E**). In summary, these results extend the antiviral spectrum of SLR14 to include a segmented negative-sense RNA virus and indicate that activating the RIG-I response, even after RVFV infection, leads to dramatic increase of survival, demonstrating the potential of SLR14 agonists as antiviral countermeasures.

**Figure 7.**
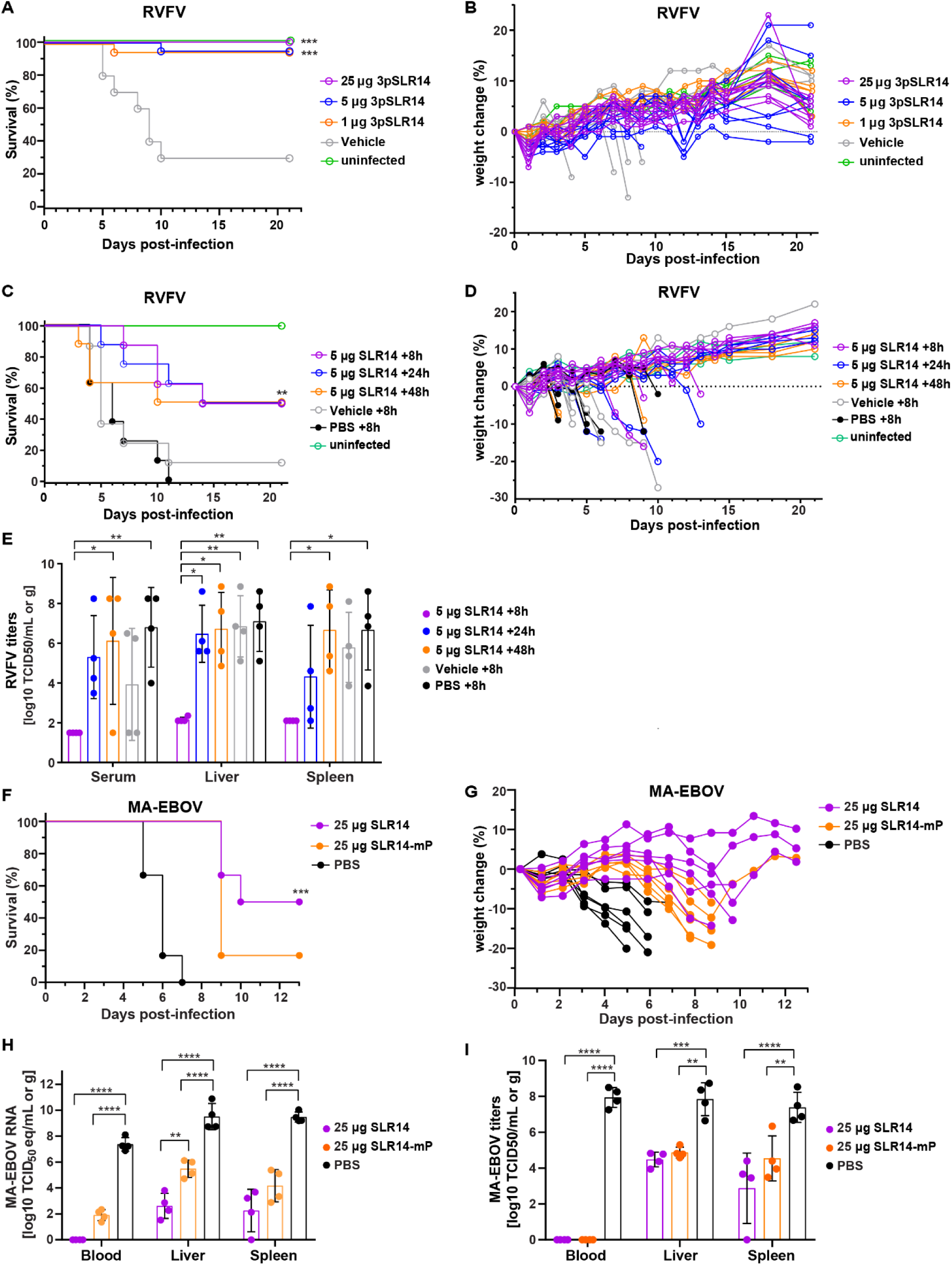
SLR14 pre-treatment protects mice against RVFV disease and EVD. (**A**) Kaplan-Meier survival plots of RVFV-infected BALB/c mice pretreated with different doses of 3pSLR14 or vehicle (jet-PEI) 6 hours prior to infection with 300 PFU RVFV (n=10 for 3pSLR14 and vehicle groups; n=4 for uninfected group). (**B**) Daily body weights normalized to day of infection and presented as percent change. Data shown for individual mice. (**C**) Kaplan-Meier survival curves of BALB/c mice infected with 300 PFU RVFV and treated with SLR14, vehicle (jet-PEI) or PBS administered IP at 8, 24, or 48 hpi (n=8 for SLR14, vehicle and PBS groups; n=4 for uninfected group). (**D**) Daily body weights normalized to day of infection and presented as percent change. Data shown for individual mice. (**E**) RVFV titers in serum, liver, and spleen at 3 dpi (n=4 per group). (**F**) Kaplan-Meier survival plots of CD-1 mice infected with 100 LD_50_ (1 FFU) of MA-EBOV and treated with 25 μg of SLR14 or SLR14-mP 8 hours later. PBS treatment was used as control (n=6 per group). (**G**) Daily body weights normalized to day of infection and presented as percent change. Data shown for individual mice. (**H**) MA-EBOV RNA levels (titer equivalents) and (**I**) MA-EBOV titers in blood, liver and spleen at 4 dpi (n=4 per group). Survival curves in (A), (C), and (F) significantly differ across groups, as determined by the Log-rank (Mantel-Cox) test (*p<*0.0001). Statistical analysis of different conditions was performed using two-way ANOVA followed by the Tukey’s post-test (E,H,I). P-values: *<0.1; **<0.01; ***<0.001; ****<0.0001; ns: not significant.

### SLR14 protects mice against EVD

We next assessed the protective effects of SLR14 against EBOV infection in the CD1 murine model. Since wild-type EBOV does not cause disease in immunocompetent mice, we used mouse-adapted EBOV (MA-EBOV), which has been shown to be uniformly lethal in this model (*71*). CD-1 mice were infected IP with a dose of 100 LD_50_ (1 FFU) and received IP treatment with SLR14, SLR14-mP or PBS 8 hours later. Mice were monitored daily for weight loss and survival for up to 28 days, and 4 mice per group were euthanized on day 4, which corresponds to peak disease, for blood and tissue sample collection to determine differences in viral loads. While all PBS-treated mice and 5 out of 6 of the SLR14-mP-treated mice succumbed to MA-EBOV infection, 50% of the mice treated with SLR14 survived to the end of the study (**Fig. 7F and G**). Pairwise comparisons revealed significantly improved survival in SLR14-treated mice compared to SLR14-mP- and PBS-treated animals (*p*=0.0054 and 0.0004, respectively), demonstrating a protective effect of SLR14 against lethal infection. At 4 dpi, both viral titers and viral RNA levels were significantly reduced in SLR14-treated mice across blood, liver, and spleen compared to PBS-treated control mice. While significantly lower viral burden was also reported in mice treated with the control SLR14-mP, the reduction of viral RNA was significantly stronger in liver and spleen tissue treated with the biologically active SLR14 (**Fig. 7H and I**), corresponding to the observed enhanced survival rates. Together, these data demonstrate that SLR14 provides potent protection against lethal MA-EBOV infection in mice when administered 8 hpi.

## Discussion

Emerging zoonotic RNA viruses are a growing public health concern, as evident from the recent SARS-CoV-2 pandemic. To strengthen preparedness for future pandemics of potentially unknown origin, it is necessary to develop novel and broad-spectrum antiviral therapeutics that can be rapidly applied in an outbreak situation. Here, we tested the antiviral efficacy of the RIG-I agonist SLR14 against five different negative-sense RNA virus species from three distinct viral families that include important human pathogens, the *Filoviridae*, *Paramyxoviridae*, and the *Phenuiviridae*, with an emphasis on the filoviruses. First, we showed that SLR14 in its biologically active 5’-terminally diphosphorylated form induces a rapid and robust IFN-I response in human primary MDMs, which are a primary site of filovirus replication. Importantly, the upregulation of most IFN genes was short-lived, which is essential to avoid contributing to persistent inflammation, while most of the assessed ISGs and cytokines were still expressed at later time points, maintaining the antiviral effect. This confirms previous observations in cultured cells treated with SLR14 or similar RIG-I activating RNAs, which showed a rapid but short-lived (24 hours or shorter) induction of IFN genes followed by prolonged ISG expression (*9, 27, 72*).

Consistent with the observed IFN-I response, SLR14 potently blocked EBOV and SUDV infection of MDMs, both when the cells were treated pre- or post-infection. Notably, we observed a modest protective effect of the controls, SLR14-OH and SLR14-mP, albeit substantially weaker than that of SLR14. This might be due to residual but inefficient RIG-I binding and activation (*24*), slightly amplifying the filovirus-induced IFN-I response. We further demonstrated the broad applicability of SLR14 by showing its protective effect against the more distantly related MARV in MDMs. Additionally, we found that a single dose of SLR14 administered 8 hpi protected 50% of mice against lethal EBOV infection. This was associated with a strong decrease in viral titers and RNA levels (**Fig. 7**). Surprisingly, we also observed a decrease in viral titers and RNA levels in SLR14-mP-treated mice. However, this was not associated with increased survival rates. These data are in line with our MDM cell culture data and suggest that SLR14-mP treatment induces a weak, inefficient antiviral response in the murine infection model that can delay EBOV replication temporarily but is eventually overcome by the infection. Importantly, viral titers and RNA levels were determined at a relatively early stages of infection (4 dpi).

The importance of RIG-I mediated IFN-I induction in controlling filovirus infection is evident from the evolution of viral proteins that antagonize this pathway. The filoviral protein VP35 blocks IFN-I induction by binding to and masking viral dsRNA, thereby preventing RIG-I activation (*73–77*), as well as further downstream by impairing the TBK1- and IKK-ε-dependent phosphorylation and activation transcription factors of IRF3/7 (*78, 79*). Indeed, mutations that disrupted the ability of VP35 to mask dsRNA resulted in virus attenuation in both cell culture and animal models (*80–82*). Despite viral interference with RIG-I signaling, filovirus infection still induces an IFN-I response. However, this response is delayed and not protective (*38, 59, 83*). Our comparative transcriptomic analysis of untreated and SLR14-treated EBOV-infected macrophages revealed similar transcriptomic signatures. However, SLR14 treatment generally enhanced the magnitude of this transcriptional response, thereby directing the host response toward protection. While SLR14 treatment amplified the expression of most of the genes also upregulated in EBOV infection alone, IFNB1 and IFNL1 upregulation was markedly lower in infected SLR14-treated samples compared to EBOV infection alone. This indicates that effective control of filovirus infection depends not simply on the induction of individual antiviral genes, but rather on a robust and well-balanced innate immune activation. As shown by RNA FISH analysis, the antiviral response appears to be initially restricted to non-infected bystander cells that lack viral proteins counteracting innate immune responses, which likely contributes to the overall lower expression levels of innate immunity-related genes in EBOV infection compared to SLR14-treated cells. This initial restriction to uninfected bystander cells may be overcome over time as the virus replicates and more PAMPs accumulate within the cell. Thus, viral interference promotes a delayed and prolonged IFN response that we also observed in this study. The crucial role of non-infected bystander cells in the induction of an innate immune response was corroborated in animal studies. Enhanced expression of the ISG MxA was detected in uninfected cells in liver samples of EBOV-infected non-human primates, whereas the MxA expression levels were considerably lower in EBOV-infected cells (*84*).

RIG-I activation induces IRF3 and NFκB signaling, which is consistent with our observation of early cytokine/chemokine expression, including IFN genes, in MDMs. Previous work on the antiviral effects of RIG-I agonists on dengue and chikungunya virus infections demonstrated the importance of an intact RIG-I/MAVS/TBK1/IRF3 axis for the protective response, whereas IFN-I signaling was largely dispensable (*32*). In contrast, 3pRNA-mediated protection against RSV infection in murine lung fibroblasts required functional IFN-I signaling (*15*). Similarly, SLR14-induced protection against SARS-CoV-2 infection in mice was dependent on IFN-I signaling, as demonstrated by the loss of protection upon IFNα/βR blockade (*28*). An intermediate protective phenotype was observed in an influenza A mouse model, in which 5′PPPRNA partially protected IFNAR-deficient mice, albeit less effectively than wild-type mice, suggesting both IFN-dependent and IFN-independent mechanisms of protection (*9*). Small-molecule activation of IRF3 was shown to reduce EBOV titers in a cell culture model despite modest IFNB induction (*85*). Our results demonstrate that SLR14 treatment elicits an early and well-controlled IFN response in EBOV-infected MDMs that might be essential for viral control. While we cannot exclude that IRF3 activation alone also contributes to the observed antiviral effects, we previously showed that IRF3 nuclear translocation in EBOV-infected macrophages failed to inhibit viral replication, suggesting that IRF3 signaling alone is insufficient (*51*). In the clinical context, an early, robust IFN-I response is generally associated with higher chances of recovery and survival, whereas delayed and prolonged IFN-I responses during later stages of infection are associated with immunopathology, driving increased severeness of disease. A small study of biomarkers in patients with EVD found that a functional IFNβ response was associated with moderate rather than severe disease (*86*), and treatment with IFNβ was shown to prolong survival of non-human primates infected with EBOV or MARV (*87*). Stimulation of RIG-I prior to EBOV infection or overexpression of the isolated RIG-I caspase activation and recruitment domains (CARDs) that mediate interaction with MAVS was shown to inhibit EBOV replication *in vitro* (*88*). Similarly, a study found that nitazoxanide enhances RIG-I and RIG-I-like receptor (RLR) signaling and can inhibit EBOV replication (*89*). These studies, in line with our data, indicate that therapeutic RIG-I activation constitutes a promising approach for the treatment of patients with EVD.

In addition to filoviruses, we describe the antiviral effect of SLR14 against the paramyxovirus NiV, which also encodes several viral proteins that interfere with innate immunity (*44, 90, 91*). In our iEC model, NiV infection led to a delayed innate immune response with minimal changes at 6 hpi and robust induction at 24 hpi. This is consistent with previous findings showing no IFN induction up to 15 hpi in NiV-infected artery iECs and significant upregulation of IFN genes and ISGs at 24 hpi compared to 8 hpi in NiV infection of human umbilical vein endothelial cells (*43, 67*). Similar to what we found for EBOV, we observed ISG expression upon NiV infection mainly in non-infected bystander cells, potentially contributing to a delayed IFN response. This is in line with previous work, suggesting that uninfected bystander cells produce ISGs in response to circulating IFN from NiV-infected cells, whereas autocrine IFN signaling is blocked in NiV-infected cells (*92*).

Delayed IFN responses, likely a consequence of viral interference with RNA sensing and RIG-I signaling, contribute to disease severity across various viral infections. This includes SARS-CoV-2 (*93*), which is sensitive to treatment with SLR14 and other RIG-I agonists (*12, 28*). Similarly, in our study, prophylactic SLR14 treatment protected mice against the phenuivirus RVFV, which antagonizes the IFN-I response via its non-structural NSs protein (*94*). SLR14 treatment up to 48 hpi still protected 50% of the animals against lethal RVFV, suggesting that RIG-I activation followed by IFN-I induction has a significant protective effect even after initial viral spread. Similar therapeutic effects of RIG-I ligands were reported for SARS-CoV-2 and influenza A virus mouse models (*9, 12, 13*). While SLR14 treatment at 24 and 48 hpi did not significantly suppress viral replication, it still achieved 50% survival, with initial mortality appearing at 3 dpi. Similarly, 5′pppRNA treatment of influenza A virus-infected mice reduced viral titers early in infection but not at later time points, despite conferring complete protection (*9*). In this study, repeated dosing of 5′pppRNA achieved additional viral titer reduction in the lung, emphasizing that optimal treatment timing is critical to control viral replication.

In conclusion, our results help explain the counter-intuitive finding that highly pathogenic viruses such as Ebola and Nipah viruses trigger robust innate and inflammatory responses yet fail to provide protection. Furthermore, our work informs the development of broad-spectrum prophylactic and therapeutic antiviral strategies based on augmenting innate immune responses. The current Bundibugyo virus outbreak, which is the second largest filovirus outbreak on record, highlights the critical importance of virus-agnostic antiviral strategies, which our work directly addresses.

## Limitations of the study

Our proof-of-concept study establishes SLR14 as a virus-agnostic therapeutic with significant potential against the most aggressive human viruses. SLR14 holds also promise as a cost-effective, large-scale manufacturable compound. However, further studies are needed to optimize SLR14 dosing regimens and tissue distribution. Our findings also support the potential of combination therapies that pair virus-agnostic immune stimulation with broad-spectrum antivirals targeting viral replication complexes, such as remdesivir, but this strategy was not evaluated in this study.

## Materials and methods

### Biosafety statement

All cell culture work with EBOV, SUDV, MARV and NiV was performed in the biosafety level 4 (BSL-4) facility of Boston University’s National Emerging Infectious Diseases Laboratories (NEIDL) following approved standard operating procedures in compliance with local and national regulations pertaining to handling BSL-4 pathogens and Select Agents.

The MA-EBOV mouse studies were performed in the BSL-4 facility of the Rocky Mountain Laboratories (RML), Division of Intramural Research, National Institute of Allergy and Infectious Diseases, National Institutes of Health. RML is an institution accredited by the Association for Assessment and Accreditation of Laboratory Animal Care International (AAALAC). Experiments were conducted in compliance with local, state and federal regulations relevant to BSL-4 operations, select agents and animal welfare.

All work with RFVF was performed in biosafety level 3 enhanced (BSL-3+) facilities at Utah State University (USU), following Select Agent-compliant operations and procedures.

### Ethical statement

RVFV mouse studies and procedures complied with USDA guidelines and were performed at AAALAC-accredited laboratory animal research facilities at USU under protocol #10248, approved by the Institutional Animal Care and Use Committee (IACUC).

The MA-EBOV mouse studies were performed following standard operating procedures approved by the RML Institutional Biosafety Committee. The work was performed in accordance with the recommendations described in the Guide for the Care and Use of Laboratory Animals of the National Institute of Health, the Office of Animal Welfare and the United States Department of Agriculture and was approved by the RML Animal Care and Use Committee (ACUC). Procedures were conducted in animals anesthetized by trained personnel under the supervision of veterinary staff. Food and water were available *ad libitum*. End point criteria as specified by RML ACUC-approved clinical score parameters were used to determine when animals were humanely euthanized.

### Cell lines

African green monkey kidney cells (Vero E6; ATCC Manassas, VA, USA; CRL-1586) were maintained in cell culture medium consisting of Dulbecco’s modified Eagle medium (DMEM; Gibco/Thermo Fisher Scientific, Waltham, MA, USA) supplemented with L-glutamine (200 mM; Thermo Fisher Scientific, Waltham, MA, USA) and 100 µg/mL primocin (Invivogen, San Diego, CA, USA). Cell culture medium was supplemented with 10% fetal bovine serum (FBS; R&D Systems, Minneapolis, MN, USA). Cells were grown at 37°C and 5% CO_2_.

### Generation of macrophages

Monocyte-derived macrophages (MDMs) were generated as described previously (*51*). Briefly, MDMs were generated from Leukoreduction system chambers (NY Biologics Inc.) using Ficoll density-gradient separation (GE Healthcare) followed by isolation of CD14+ monocytes by magnetic bead selection (Milteny Biotech). CD14-selected cells were seeded into ultra-low attachment 6-well plates (Corning) for differentiation into MDMs using RPMI medium supplemented with penicillin (50 units/mL), streptomycin (50 mg/mL), HEPES (10 mM), 5% FBS (certified One Shot, Gibco) supplemented with 100 ng/mL human GMCSF (Peprotech) for 6–8 days. Differentiated MDMs were lifted from plates using Cell Stripper (Corning), and 3x10^5^ cells were seeded into 24-well culture plates or 7x10^4^ cells into Lab-Tek II 8-well chamber slides (Nunc/Thermo Fisher). To account for donor variations, experiments were performed with cells obtained from at least three different donors as noted in the figure legends. Throughout the figures, each donor is represented by a donor-specific symbol.

### Derivation of human iECs from induced pluripotent stem cells

Human induced pluripotent stem cells (iPSCs) from previously published clone BU1 (available from crem.bu.edu) were used for these studies and maintained in mTesr1 medium (StemCell Technologies). All iPSC related work was performed under regulatory approval of Boston University’s Institutional Review Board. For passaging, cells were dissociated for 3-5 minutes using Gentle Cell Dissociation Reagent (StemCell Technologies) and then manually lifted and gently pipetted into a new well. For endothelial cell (iEC) differentiation, iPSCs were dissociated to single cells using Gentle Cell Dissociation Reagent for 15-20 minutes and replated at a density of 1x10^6^ cells per well of a 6-well plate in mTesr1 with ROCK inhibitor. The next day, cells were treated with BMP4 (25 ng/mL), FGF2 (50 ng/mL), and CHIR-99021 (6 μM) for 24 hours in STEMdiff APEL 2 media at a volume of 1 mL per well. Cells were then treated with recombinant human BMP4 (25 ng/mL) and FGF2 (50 ng/mL) in STEMdiff APEL2 media for 24 hours (2 mL/well). Endothelial specification was then initiated through treatment with VEGFA (200 ng/mL) and DAPT (10 μM) in STEMdiff APEL2 medium for 72 hours with medium (2 mL/well) replaced daily. iECs were then purified by magnetic activated cell separation using microbeads targeting hCD31 and hCD144. Purified iECs were grown and expanded in EGM2 medium. Differentiated iECs were seeded at 2x10^5^ into 24-well culture plates or 1.5x10^5^ cells into Lab-Tek II 8-well chamber slides (Nunc/Thermo Fisher).

### Viruses

EBOV Mayinga (Genbank NC_002549.1), SUDV Gulu (GenBank number NC_006432.1) and NiV Bangladesh (GenBank number AY988601.1) were kindly provided by Heinz Feldmann, NIH NIAID RML. Recombinant EBOV expressing ZsGreen (EBOV-ZsG; GenBank number OL956940.1) was previously described (*65*). Virus stocks were propagated in Vero E6 cells in cell culture medium supplemented with 2% FBS and purified by ultracentrifugation through a 20% sucrose cushion as previously described (*51*). Virus titers were determined in Vero E6 cells by tissue culture infectious dose 50 (TCID_50_) assay using the Spearman and Kärber algorithm (*95*). MA-EBOV was propagated in Vero E6 cells, sequence confirmed and stored in liquid nitrogen as previously described (*71*). Virus titers were determined in Vero E6 cells by focus-forming unit assay.

The molecular clone of RVFV, strain ZH501 (*96*), was kindly provided by Dr. Stuart Nichol (CDC, Atlanta, GA). The virus stock (rescued in BSRT7 cells and passaged 3 times in Vero E6 cells) was verified by Next-Generation Sequencing and the concentration of infectious virus determined by standard plaque assay.

### Preparation of SLR14, p3SLR14, SLR14-mP and SLR14-OH

Di- and triphosphorylated SLR14 RNAs (SLR14 and p3SLR14) were synthesized, deprotected and purified as previously described (*21, 26*). SLR14-mP and SLR14-OH were prepared as described in (*97*). The molecular mass of SLR14 is 10,376 Da.

### Viral infections of MDMs and iECs

MDMs or iECs seeded in 24-well plates or Lab-Tek II 8-well chamber slides at the cell numbers indicted above were infected with sucrose cushion-purified virus stocks. Prior to infection the virus stocks were diluted in the appropriate cell culture medium for MDMs or iECs to obtain the desired MOI. Cell supernatants (with or without SLR14 derivates) were removed, and 0.5 mL (24-well format) or 0.3 mL (chamber slides) of virus inoculum was added. The cells were incubated until time of harvest. Cell culture medium was not changed or added to avoid disturbance of the primary cells.

### SLR14 treatment of MDMs and iECs

SLR14 and SLR14 derivates were diluted to 100 ng/μL in sterile H_2_O and stored in aliquots at - 80°C to avoid freeze-thaw cycles. Except for the initial testing of optimal SLR14 concentrations, 1 µg of SLR14 (or SLR14 derivates) was used for MDMs per sample and 25 ng for iECs. SLR14 was added to 50 μL of Opti-MEM (Thermo Fisher) and mixed with TransitX2 transfection reagent (Mirus) at a ratio of 1.5 μL TransitX2 per 100 ng of SLR14. The solution was incubated for 15-30 minutes at room temperature and then added onto cells seeded in 24-well plates. Cells were incubated with SLR14 for the indicated times. For SLR14 pre-treatment, The SLR14 transfection solution was removed from the cells after 6 hours and virus inoculum was added. For SLR14 post-infection treatment, the SLR14 transfection solution was pipetted into the cell culture medium of the infected cells at the indicated time points.

### Preparation of RNA samples and RT-qPCR analysis

Total RNA used for RT-qPCR was isolated at the indicated time points from 24-well plates (MDMs or iECs) using TRIzol Reagent (Invitrogen) according to the manufacturer’s protocol and BSL-4 inactivation SOPs. RNA concentration was determined using the NanoDrop 1000 Spectrophotometer (Thermo Scientific). 5 ng of RNA was analyzed by quantitative reverse transcription PCR (RT-qPCR) using the QuantiNova SYBR green RT-PCR kit or (QIAGEN) and validated QuantiTect primer assays for IFNB, TNFA, CXCL10, ISG15, OASL andCCL5 (QIAGEN). Validated QuantiTect primer assays (QIAGEN) for β-2-microglobulin were used as an endogenous control for normalization. Fold expression levels compared to control samples collected at each corresponding time point were quantified using the ΔΔCT method (Biorad CFX Manager 1.5 software). RT-qPCR of viral genes was performed using primers binding to the EBOV VP35 gene (GAGGAAAATTTTGGGAAACC; GCCTGGAACTCAGCATG) or the NiV N gene (GAGGAATTGATCAGAACATGG; GATATCTTGATCACTGCCTCC).

### Preparation of RNA samples and RNA-Seq analysis

RNA used for RNA-Seq was isolated at the indicated time points from 24-well plates (MDMs or iECs) using TRIzol Reagent (Invitrogen) and processed up to the phase separation step according to the manufacturer’s protocol and BSL-4 inactivation SOPs. The aqueous phase containing RNA was then purified using the RNeasy Micro kit (QIAGEN) according to the manufacturer’s protocol including DNase I digestion and optional wash steps. RNA-Seq analysis was performed by Boston University Chobanian & Avedisian School of Medicine’s Microarray and Sequencing Resource Core Facility. RNA concentration and quality was assessed using a Bioanalyzer. RNA-Seq libraries were prepared using poly(A) selection to enrich for polyadenylated mRNA. Libraries were sequenced on an Illumina NextSeq 2000 platform using a P4 flow cell (2 × 50 bp paired-end reads), generating approximately 1.7 billion total reads per run.

### Bioinformatic analysis of RNA-Seq data

The quality of the raw data was assessed using FastQC v.0.11.7 (*98*). The sequence reads were aligned to the GRCh38 reference with added sequences of EBOV, SUDV and NiV Bangladesh using STAR v.2.6.0 (*99*). Counts per gene were summarized using the featureCounts function from the subread package v.2.0.3 (*100*). The edgeR package v.4.2.0 (*101*) was used to import, organize, filter and normalize the counts, and the matrix of counts per gene per sample was then analyzed using the limma/voom normalization method (*102*). Genes were filtered based on the standard edgeR filtration method using the default parameters for the “filterByExpr” function, which excludes genes with low expression across all samples. Specifically, genes with fewer than 10 counts were removed from the dataset prior to differential expression analysis. After Principal Component Analysis (PCA), differential expression testing for the samples was done in comparison to appropriate controls at each time point. The limma package v.3.60.0 (*102*) with its voom method, namely, linear modelling and empirical Bayes moderation was used to test differential expression (moderate t-test). P-values were adjusted for multiple testing using Benjamini-Hochberg correction (false discovery rate-adjusted p-value; FDR). Differentially expressed genes for each comparison were visualized using Glimma v.2.14.0 (*103*), and FDR < 0.05 was set as the threshold for determining significant differential gene expression. Functional predictions were performed using the fgsea v.1.30.0 package (*104*) for gene set analysis. Genes with the greatest log2 fold changed in SLR14 only conditions were selected by 7 log2 fold change > and FDR < 0.05. The selected genes were compared to the gene expression values of untreated and treated samples of EBOV and SUDV.

### Immunofluorescence

Infected MDMs or iECs seeded in 8-well chamber slides were fixed with 10% formalin for at least 6 hours and removed from BSL-4 in accordance with approved inactivation SOPs. The fixed cells were permeabilized with 0.1% Triton X100 (Boston Bioproducts, Milford, MA, USA) for 10 minutes at room temperature, incubated in 0.1 M glycine (Boston Bioproducts, Milford, MA, USA) for 5 min at room temperature, and subsequently incubated in blocking buffer (2% bovine serum albumin, 0.2% Tween 20, 3% glycerol, and 0.05% sodium azide in PBS) or goat serum (Jackson Immunoresearch) for 20-60 minutes at room temperature. After each step, the cells were washed three times in PBS. Cells were incubated overnight at 4°C with goat anti-EBOV VP35 (Antagene, custom-made), rabbit anti-SUDV NP (IBT), rabbit-anti-MARV nucleocapsid (anti-MARV NC; Mühlberger laboratory) or rabbit-anti NiV-N (GeneTex) antibodies. The cells were washed four times in PBS and incubated with an Alexa Fluor AF488 conjugated anti-goat or anti-rabbit secondary antibody and 4’,6-diamidino-2-phenylindole (DAPI; Sigma-Aldrich at 200 ng/mL for nuclei staining) for 1 hour at room temperature. Cells infected with EBOV-ZsG were only stained with DAPI in PBS for 20 minutes without prior permeabilization after formalin fixation.

Images were acquired using a Nikon TS100 Eclipse microscope and Nikon DS Qi1Mc camera with NIS Elements F software or with a Nikon Eclipse Ti2 microscope with Photometrics Prime BSI camera and NIS Elements AR software. Fiji Image J software (https://github.com/fiji) was used to create merge channel images and add scale bars. Except for EBOV-ZsGreen quantification, where 2 images per condition were quantified, at least 5 images per experiment and donor were quantified using the QuPath cell detection feature (https://qupath.github.io/) to identify nuclei by DAPI staining. Infection rates were quantified using mean fluorescence signal for the GFP channel (*105*). Percentage for each subset of cells was graphed in Prism.

### RNA fluorescence in situ hybridization analysis

For RNA fluorescent in situ hybridization (FISH) analysis, infected MDMs or iECs seeded in 8-well chamber slides were fixed with 10% formalin for at least 6 hours and removed from the BSL-4 laboratory in accordance with approved inactivation SOPs. RNA FISH was performed using the RNAscope Multiplex Fluorescent V2 kit (Advanced Cell Diagnostics/BioTechne). EBOV and NiV RNA was detected using custom-designed probes (Advanced Cell Diagnostics/BioTechne). Specifically, viral genomic RNA for EBOV using probes targeting the negative-sense genomic sequences of the NP gene or viral mRNA for NiV targeting the L gene transcript were detected in channel 2 and stained with Opal 520 fluorophore (Perkin-Elmer). Cellular ISG15 mRNA was detected in using a probe in channel 1 (Advanced Cell Diagnostics/BioTechne) and stained with Opal 690 fluorophore (Perkin-Elmer). All FISH probe sets were designed by Advanced Cell Diagnostics and the sequences of these probe sets are proprietary. Staining was performed according to the manufacturer’s protocol for adherent cell samples, with the exception of an additional HRP blocking step following the signal development of the probes detecting viral RNA as per the manufacturer’s recommendation. Nuclei were stained with kit-supplied DAPI following the manufacturer’s protocol. Coverslips were mounted on slides using FluorSave mounting medium, and slides were subsequently stored at 4°C prior to imaging. Images were acquired using a Nikon Ti2 Eclipse microscope and Photometrics Prime BSI camera with NIS Elements AR software. Fiji Image J software (https://github.com/fiji) was used to create merge channel images and add scale bars. Per experiment and donor at least 6 images were quantified using the QuPath cell detection feature (https://qupath.github.io/) to identify nuclei by DAPI staining, infection and ISG15 staining were identified using a composite classifier based on mean fluorescence signal for each channel (*105*). Percentage for each subset of cells was graphed in Prism.

### IFNα ELISA

The IFNα levels in the serum samples of SLR14-treated mice were quantitated using the VeriKine-HS Mouse Interferon-Alpha All Subtype ELISA Kit (PBL Assay Science) according to the manufacturer’s instructions.

### Infection of mice with RVFV

Male and female 18–20 g BALB/c mice were obtained from Charles River Laboratories, acclimated for 7 days prior to the start of RVFV experiments, and fed Harlan Lab Block and tap water *ad libitum*.

Equal numbers of male and female BALB/c mice were weighed prior to each study and sorted into cohorts (*n*=8-10 per group) to minimize weight differences across experimental groups. Immediately prior to dosing, the SLR14 derivates and PEI (Polyplus Transfection) were separately diluted in 5% glucose and briefly vortexed. SLR14 and PEI dilutions were then combined and administered by IP injection of 0.1 mL per mouse. Control groups were treated with an equal volume of PEI solution in 5% glucose or PBS. RVFV was diluted in sterile Minimal Essential Medium and inoculated by subcutaneous injection of 0.1 mL containing 300 PFU of RVFV. Animals were observed for 21 days for morbidity (weight loss) and mortality. While the post-exposure experiment was conducted with diphosphorylated SLR14 (termed SLR14 in this manuscript), limitations in the availability of material necessitated the use of triphosphorylated SLR14 (3pSLR14) for the pre-exposure RVFV study. As described in the introduction, triphosphorylated and diphosphorylated dsRNA ligands can be used interchangeably as they both have similar high potency, which has been well established for the SLR14 family of agonists (*21, 22, 24*).

RVFV titers were determined by TCID_50_ assay on Vero 76 cells as previously described (*106*) and calculated by the Reed-Muench method (*107*). The lower assay limits of detection were 1.49 log_10_ TCID_50_/mL of serum and 2.1 log_10_ TCID_50_/g of tissue, and the upper limit of detection was 8.85 log_10_ TCID_50_/g of tissue. In samples presenting with virus outside of the limits of detection, a value representative of the lower or upper limit was assigned for statistical analysis.

### Infection of mice with EBOV

Female CD-1 (ICR) mice (6 weeks old, n=10 per group) were obtained from Inotiv and infected with 100 LD_50_ (1 focus-forming unit, FFU) of MA-EBOV in 100 µL total volume intraperitoneally (IP) at 2 sites in the lower abdomen on day 0 as previously described (*71*). Treatment with SLR14 and SLR14-mP (25 µg per mouse combined with *in vivo*-jetPEI transfection reagent at an N/P ratio of 8 according to the manufacturer’s instructions) was delivered as a single injection IP in a volume of 100 µL 8 hpi. Control mice received 100 µL of sterile PBS. At 4 dpi, 4 mice per group were euthanized and blood (EDTA), liver and spleen samples were collected for virologic analysis. Viral RNA levels and virus titers were determined as previously described (*108*). All remaining mice (n=6 per group) were observed daily post-infection for signs of disease and euthanized when IACUC-approved endpoint criteria were reached.

### Statistical analysis

Statistical analysis was performed using GraphPad Prism 7. P values were calculated using one-way ANOVA followed by Dunnett’s multiple-comparisons test, two-way ANOVA followed by Tukey’s multiple-comparisons test, or an unpaired, two-tailed Student’s *t*-test, as appropriate. Log-rank (Mantel-Cox) test were used for Kaplan-Meier survival curve comparisons. Symbols represent, *: p < 0.05; **: p < 0.01; ***: p < 0.001; ****: p < 0.0001; ns: not significant (p ≥ 0.05).

## Acknowledgements

The authors are grateful to Mitchell White and Dylan Deeney, Boston University and Sarah Fergione, Yale University, for technical assistance. We would like to thank the Boston University Microarray & Sequencing Resource Core for excellent sequencing services. B.G. thanks Dr. Susan Sobolov, of RIGImmune for the gift of diphosphorylated SLR14 used to conduct the post-exposure studies on RVFV. This work was supported by the Howard Hughes Medical Institute Emerging Pathogens Initiative (Agreement dated 9/16/22; Lead Investigator A.M.P.; Co-investigators E.M. and G.M.), Gates Foundation grant INV-048926 (E.M., D.N.K. and G.M.), National Institutes of Health grants R01HL095993 (D.N.K.) and P01HL170952 (D.N.K.), NHLBI grant 5K08HL173561 (A.M.H.), and the Emerging Respiratory Pathogen Award from the American Lung Association (D.N.K. and E.M.). A.M.P. is an Investigator in the Howard Hughes Medical Institute. The RVFV studies were supported by the pre-clinical services program offered through the National Institute of Allergy and Infectious Diseases, National Institutes of Health (Contract No. HHSN272201700041I). We are also grateful to all members of the Rocky Mountain Veterinary Branch, NIAID, NIH, particularly the animal caretakers, for their support of this study. This research was supported by the Intramural Research Program of the NIH (ZIA AI001254 to AM). The contributions of the NIH authors are considered works of the United States Government. The findings and conclusions presented in this paper are those of the authors and do not necessarily reflect the views of the NIH or the U.S. Department of Health and Human Services. The funders had no role in study design, data collection and interpretation, or the decision to submit the work for publication.

## Supplementary data

**Table S1.** IFNɑ serum concentrations 6 hours after SLR14 treatment of BALB/c mice. IFNɑ serum concentrations were quantified by ELISA. Each row represents one animal. Since the measured IFNα concentrations were outside the range of the standard curve, a sigmoidal, 4PL model was fitted to the standard curve data resulting in an R^2^ of 0.9934 using GraphPad Prism version 11 for Mac to interpolate the IFNα concentrations.

| Group | Animal ID | Sex | SLR14 dose (µg) | Treatment Route | IFNα concentration (pg/ml) |
| --- | --- | --- | --- | --- | --- |
| 1 | 630 | M | 25 | IV | 18,331 |
| 1 | 636 | M | 25 | IV | 17,453 |
| 1 | 641 | F | 25 | IV | 17,555 |
| 1 | 642 | F | 25 | IV | 15,851 |
| 2 | 575 | M | 25 | IP | 3,736 |
| 2 | 633 | M | 25 | IP | 3,531 |
| 2 | 644 | F | 25 | IP | 3,759 |
| 2 | 645 | F | 25 | IP | 3,763 |
| 3 | 632 | M | 50 | IP | 14,327 |
| 3 | 637 | M | 50 | IP | 16,750 |
| 3 | 640 | F | 50 | IP | 11,632 |
| 3 | 643 | F | 50 | IP | 16,436 |
| 4 | 634 | M | vehicle | IP | <2.38 |
| 4 | 635 | M | vehicle | IP | <2.38 |
| 4 | 638 | F | vehicle | IP | <2.38 |
| 4 | 639 | F | vehicle | IP | <2.38 |

**Figure S1.**
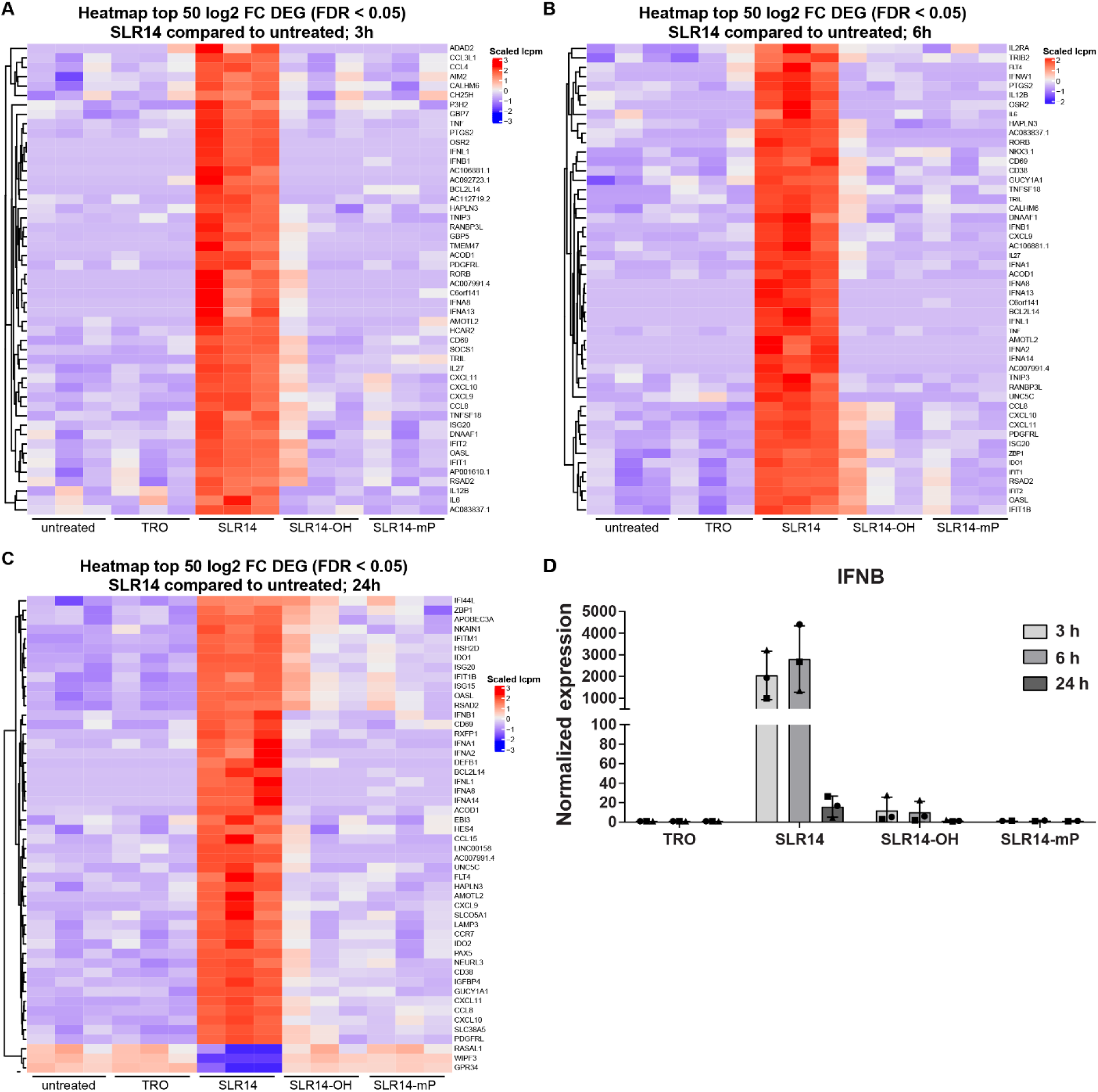
SLR14 treatment of MDMs induces a strong interferon and inflammatory response. MDMs from three different donors seeded in 24-wells were transfected with 1 μg SLR14, 1 μg SLR14-mP, 1 μg SLR14-OH, transfection reagent only (TRO), or left untreated. Total cellular RNA was harvested at the indicated time points for RNA-Seq analysis. (**A-C**) Heatmap of top 50 log2 FC genes of SLR14-treated MDMs compared to untreated cells at **(A)** 3 hours, **(B)** 6 hours, and **(C)** 24 hours. Replicates represent the three donors. **(D)** Quantification of IFNB RNA by RT-qPCR over time. Beta-2-microglobulin primers were used for normalization.

**Figure S2.**
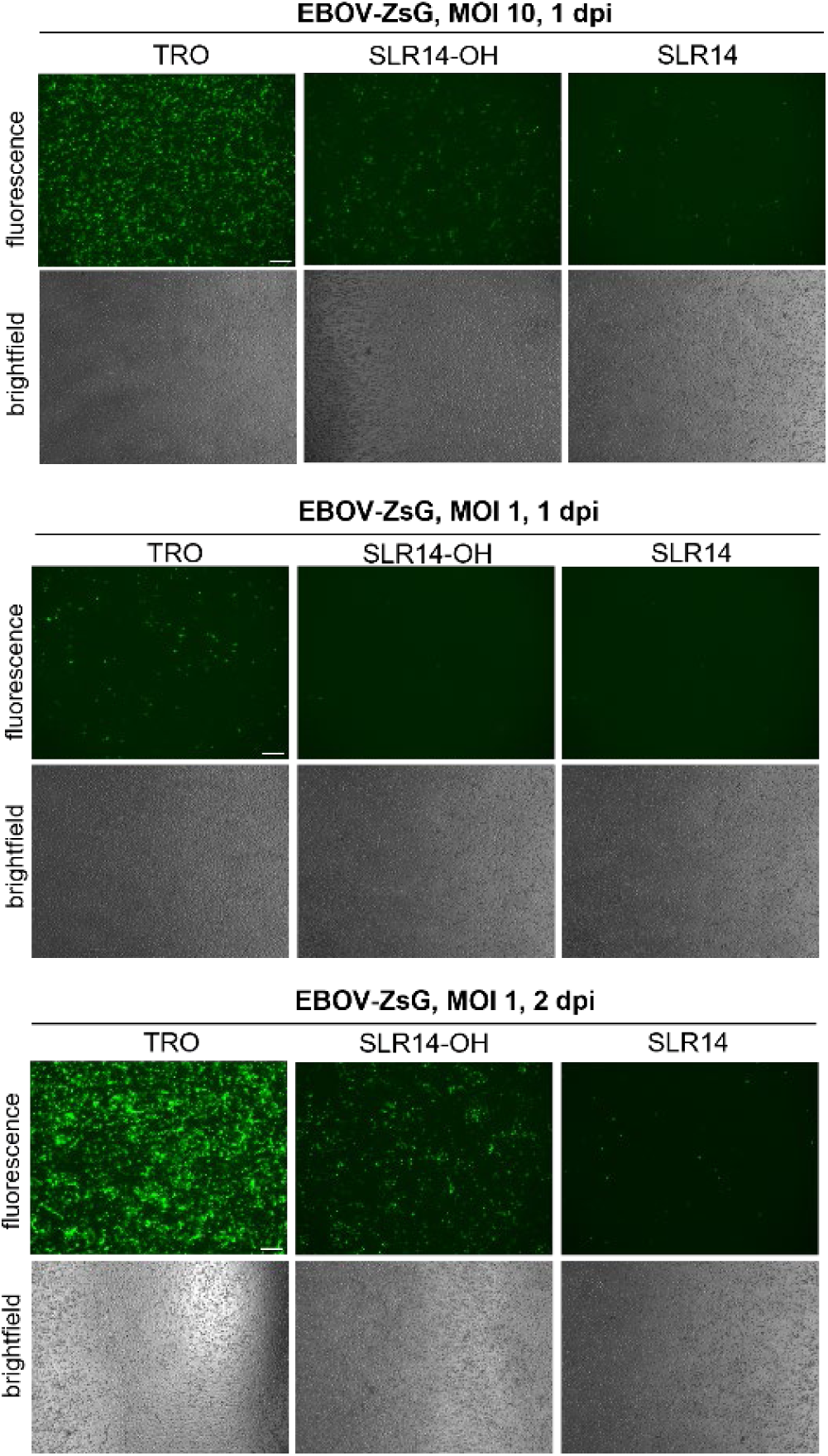
Prophylactic treatment of MDMs with SLR14 inhibits EBOV-ZsGreen infection. MDMs from three different donors seeded in 24-wells (3x10^5^/well) were transfected with 1 μg SLR14, 1 μg SLR14-mP, 1 μg SLR14-OH, transfection reagent only (TRO), or left untreated. 6 hours post-transfection, cells were infected with recombinant EBOV expressing ZsGreen (EBOV-ZsG) at MOI 1 or 10 as indicated and imaged at 1 and 2 dpi. Representative images are shown. Scale bars = 250 µm.

**Figure S3.**
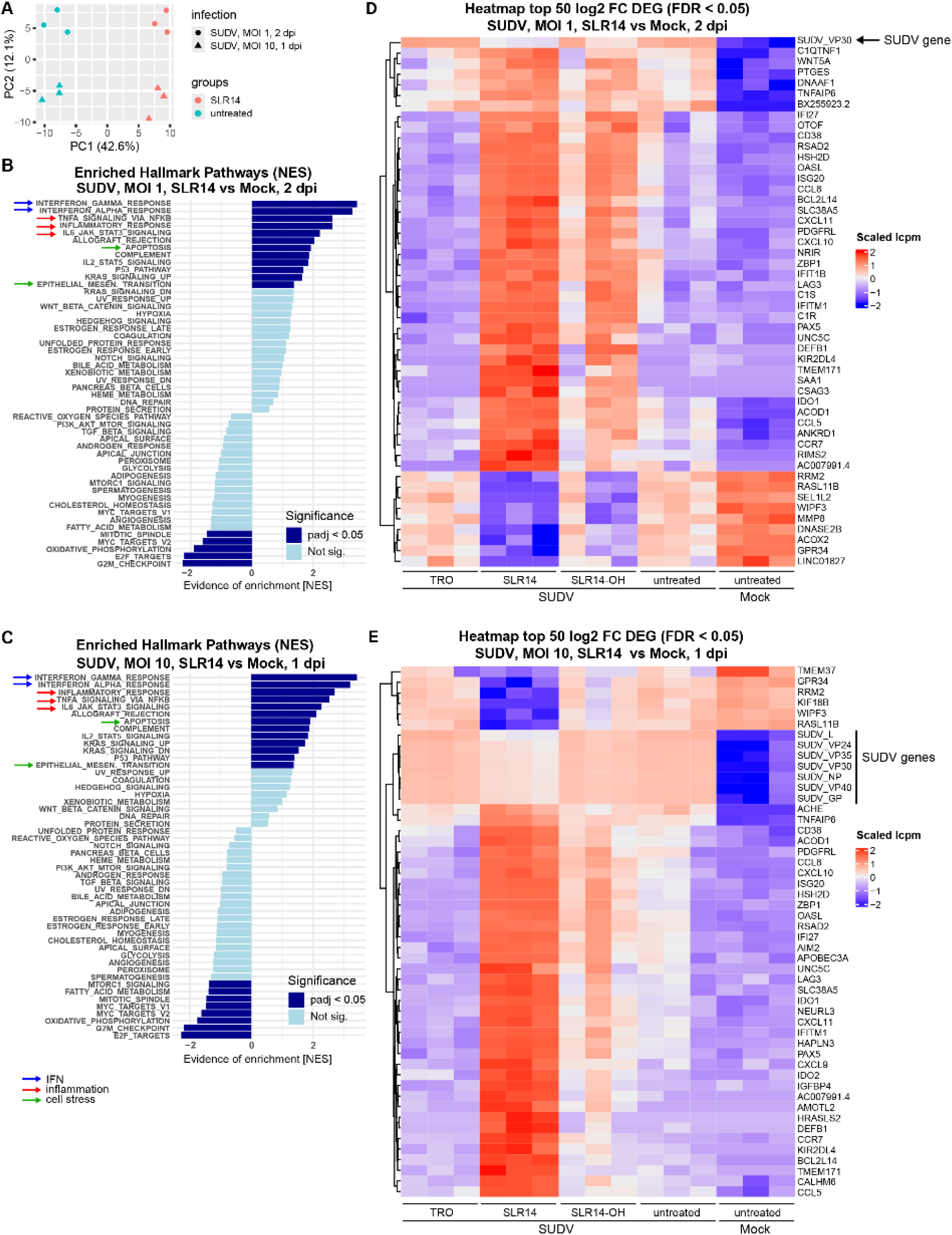
SLR14 treatment induces IFN and inflammatory responses in SUDV-infected MDMs. MDMs from three different donors seeded in 24-wells (3x10^5^/well) were transfected with 1 μg SLR14, 1 μg SLR14-OH, transfection reagent only (TRO), or left untreated. 6 hours post-transfection, the cells were infected with SUDV at MOI 1 or MOI 10 and harvested at 1 or 2 dpi. Total cellular RNA was isolated and used for bulk RNA-Seq analysis. (**A**) PCA of the transcriptomic response of the t respective treatments. Different SUDV MOI marked by symbols, different treatment conditions marked by colors. Each sample generated in 3 donors. (**B, C**) GSEA using the Hallmark gene sets on the top DEG sets in SLR14-treated, SUDV-infected MDMs compared to untreated, mock-infected controls. **(B)** SUDV MOI 1, 2d; **(C)** SUDV MOI 10, 1d. Light blue bars, not significant; dark blue bars, statistically significant (p < 0.05). (**D, E**) Heatmap of top 50 log2 FC genes in SLR14-treated, SUDV-infected MDMs compared to mock-infected controls for all tested treatment conditions. Replicates represent the three donors. **(D)** SUDV MOI 1, 2d; **(E)** SUDV MOI 10, 1d.

**Figure S4.**
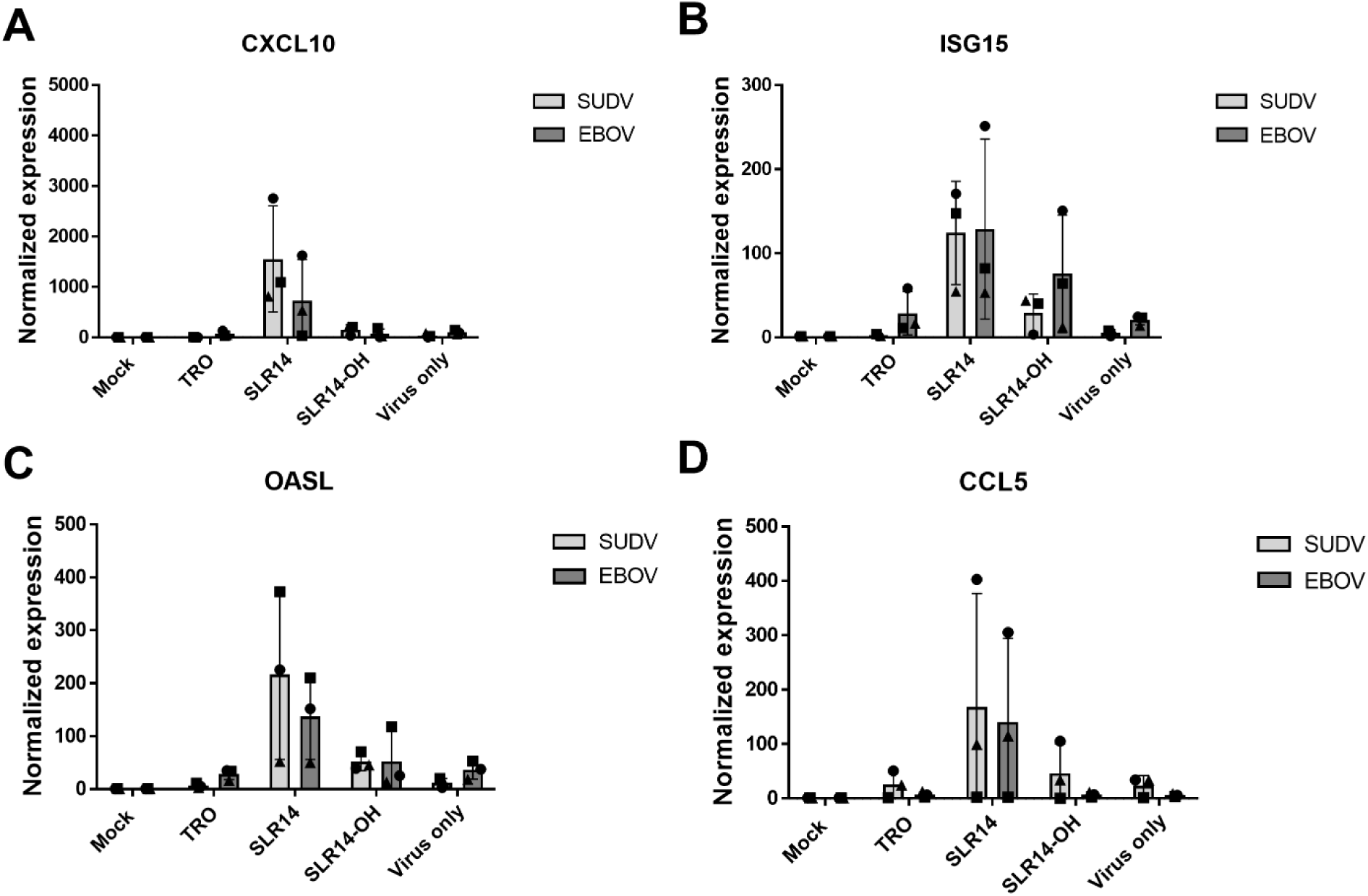
ISG and chemokine expression in SLR14-treated and EBOV or SUDV infected MDMs. MDMs from three different donors seeded in 24-wells (3x10^5^/well) were transfected with 1 μg SLR14, 1 μg SLR14-OH, transfection reagent only (TRO), or left untreated. 6 hours post-transfection, the cells were infected with EBOV or SUDV at MOI 1 and harvested at 2 dpi. Total cellular RNA was isolated at indicated time points. Quantification of **(A)** CXCL10, **(B)** ISG15, **(C)** OASL, **(D)** CCL5 RNA by RT-qPCR. Beta-2-microglobulin primers were used for normalization. Data presented as fold expression relative to mock control. Symbols indicate different donors.

**Fig. S5.**
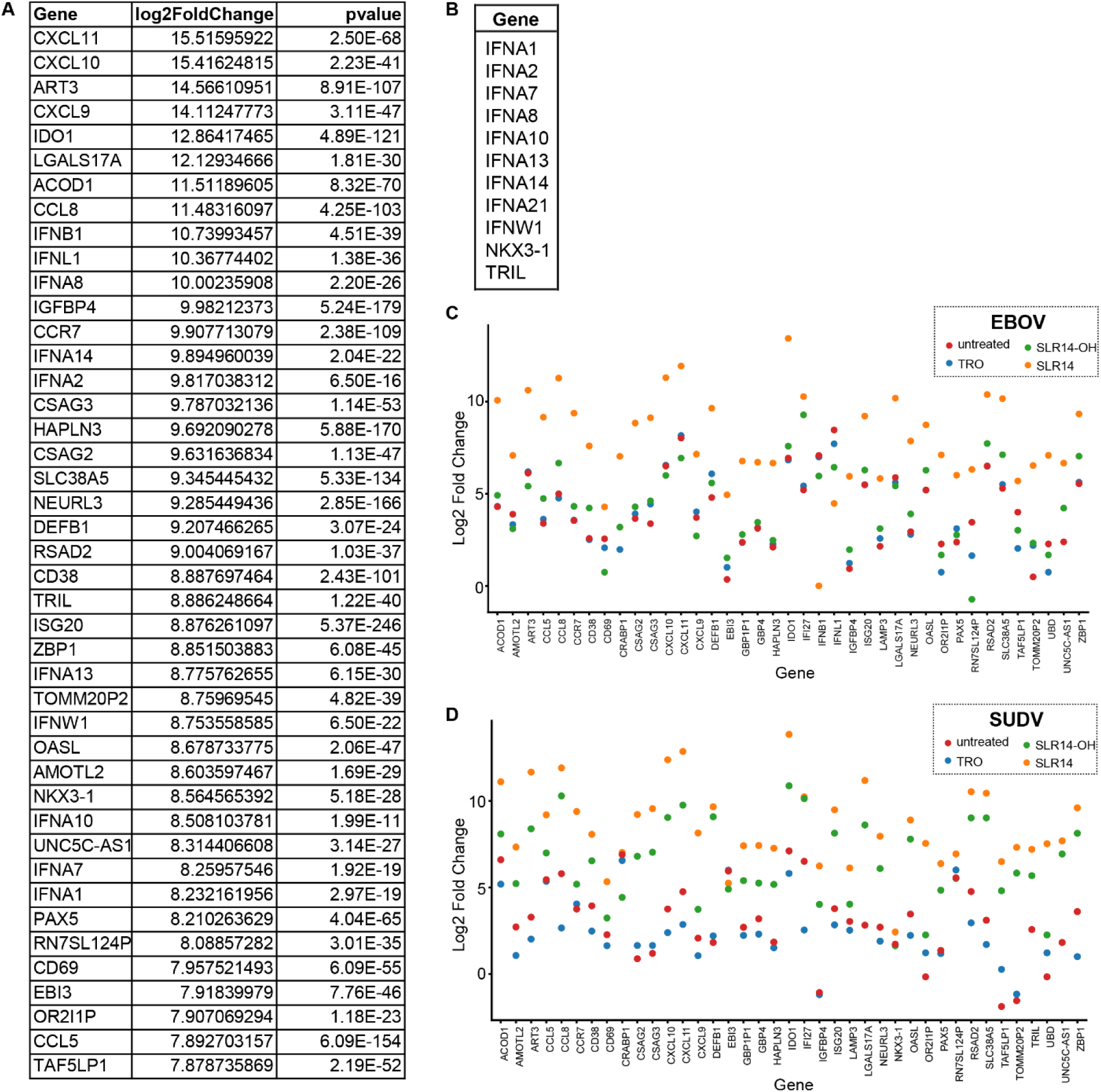
Supplemental figure to figure 4. (**A**) List of the 50 most highly upregulated genes in SLR14-treated, uninfected MDMs at 24 hours post-treatment. (**B**) List of the 11 genes that were exclusively upregulated in SLR14-treated, uninfected MDMs at 24 hours post-treatment. (**C and D**) Comparison of the 39 shared genes upregulated by SLR14 treatment in uninfected cells to the same set of genes in (**C**) EBOV and (**D**) SUDV infection under various treatment conditions. Treatment conditions are visualized by different colors.

**Figure S6.**
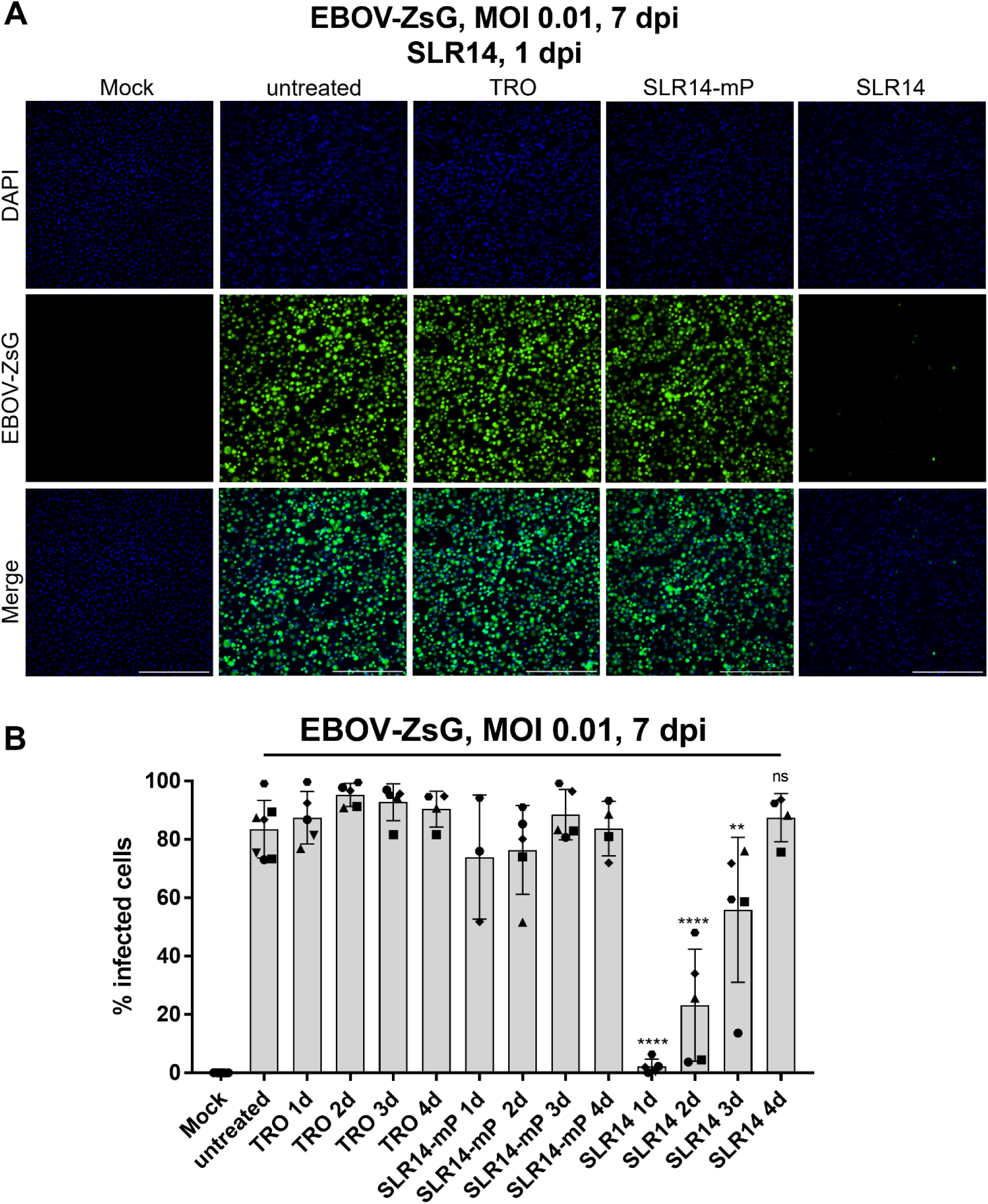
SLR14 blocks EBOV-ZsGreen infection of MDMs when administered post-infection. MDMs obtained from 3 to 5 donors seeded in 24-well plates (3x10^5^/well) were mock-infected or infected with EBOV-ZsG at MOI 0.01. At 1 to 4 dpi, the cells were transfected with 1 μg SLR14, 1 μg SLR14-mP or transfection reagent only (TRO), or left untreated. Cells were imaged at 7 dpi. (**A**) Representative images of samples treated at 1d with SLR14. Scale bars = 500 µm. (**B**) EBOV-ZsG infection rates as quantified from multiple images (n ≥ 2 per donor) of MDMs using QuPath. Donors represented by different symbols. Significant differences between conditions compared to untreated control were determined by unpaired two-tailed T-test, P-values: *<0.1; **<0.01; ***<0.001; ****<0.0001; ns: not significant.

**Figure S7.**
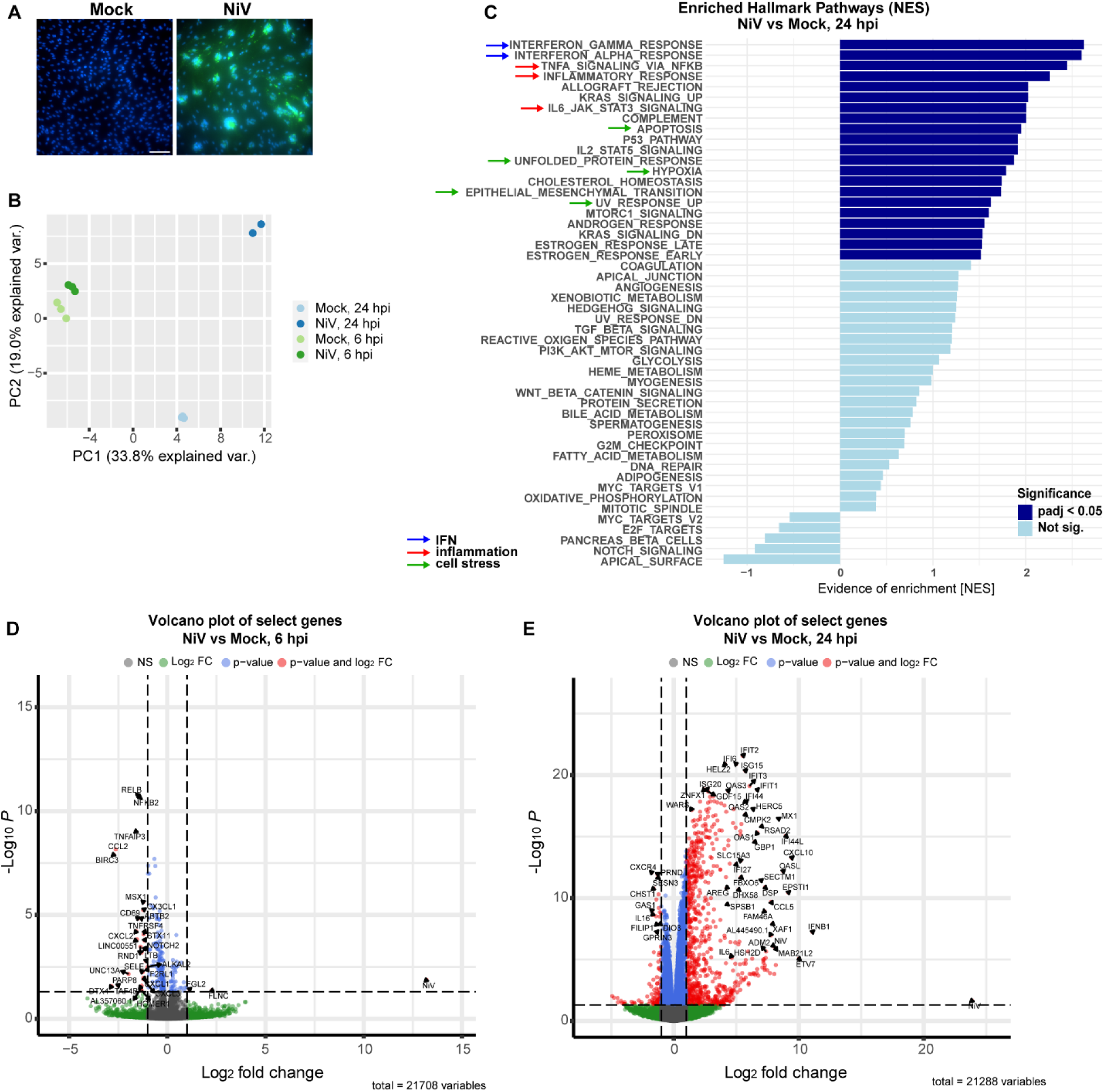
Inflammatory responses in iECs infected with NiV. **(A)** 1.5x10^5^ iECs seeded per well of 8-well chamber slides were infected with NiV at and MOI of 10. Scale bar = 100 µm **(B)** iEC seeded per well of 24-well plates were infected with NiV Bangladesh at MOI 10. The cells were harvested at the indicated time points and subjected to RNA-Seq. PCA of the transcriptomic response of the respective treatments. Each sample generated in triplicate except NiV, 24 h which was done in duplicate. **(C)** GSEA using the Hallmark gene sets on the top DEG sets in NIV-infected iECs compared to untreated, mock-infected controls at 24 hpi. **(D, E)** Volcano plot of DEGs in NiV-infected MDMs compared to mock-infected controls. Select IFNs, ISGs, and viral genes are labeled. **(D)** NIV, 6 hpi. **(E)** NIV, 24 hpi.

**Figure S8.**
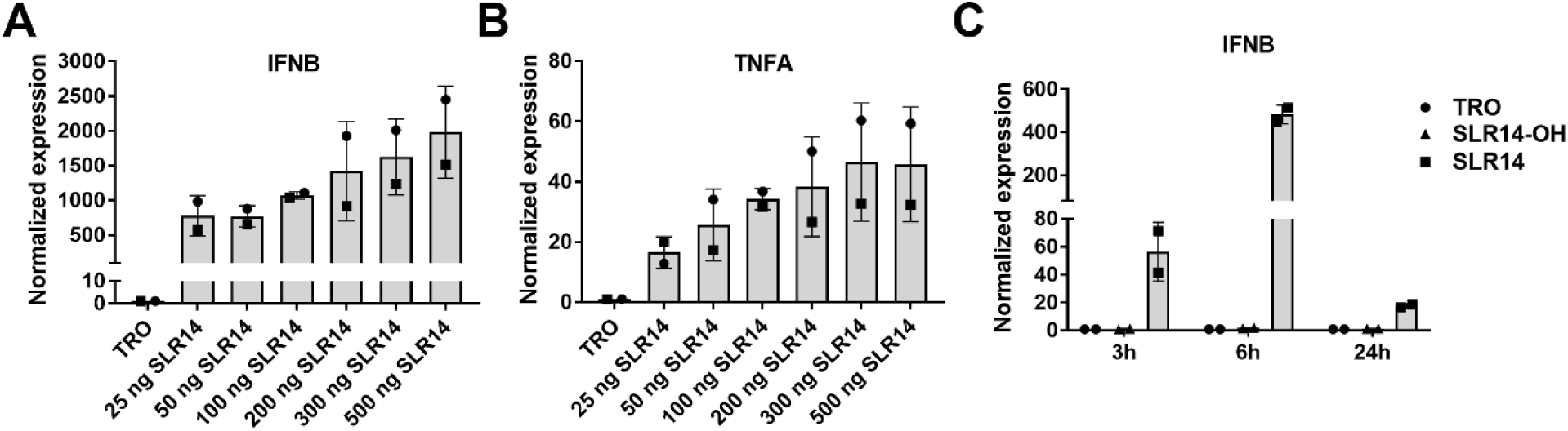
Inflammatory responses in iECs infected treated with SLR14. (**A and B**) iECs seeded in 24-well plates (1.8x10^5^/well) were transfected with increasing amounts of SLR14 or transfection reagent only (TRO) as indicated. 6 hours post-transfection, total RNA was harvested and used for RT-qPCR using primers binding to (**A**) IFNB1 or (**B**) TNFA mRNA. Normalization of RT-qPCR data was performed using beta-microglobulin as the reference gene. (**C**) iECs were transfected with 25 ng SLR14, 25 ng SLR14-OH, or TRO. Total cellular RNA was harvested at the indicated time points and subjected to RT-qPCR analysis using IFNB1-specific primers. Data presented as fold expression relative to TRO control. Different symbols represent two biological replicates.

## References

1. J. W. Schoggins, Interferon-Stimulated Genes: What Do They All Do? Annu Rev Virol 6, 567–584 (2019).

2. E. V. Mesev, R. A. LeDesma, A. Ploss, Decoding type I and III interferon signalling during viral infection. Nat Microbiol 4, 914–924 (2019).

3. S. Iurescia, D. Fioretti, M. Rinaldi, The Innate Immune Signalling Pathways: Turning RIG-I Sensor Activation Against Cancer. Cancers (Basel) 12, (2020).

4. Y. Jiang, H. Zhang, J. Wang, J. Chen, Z. Guo, Y. Liu, H. Hua, Exploiting RIG-I-like receptor pathway for cancer immunotherapy. J Hematol Oncol 16, 8 (2023).

5. G. Mandracci, N. Soliman, N. El Khawanky, Overcoming Immune Therapy Resistance in Cancer Through Innate Immune Reprogramming. Int J Mol Sci 26, (2025).

6. R. C. Ireton, M. Gale, Jr., RIG-I like receptors in antiviral immunity and therapeutic applications. Viruses 3, 906–919 (2011).

7. H. Y. Yong, D. Luo, RIG-I-Like Receptors as Novel Targets for Pan-Antivirals and Vaccine Adjuvants Against Emerging and Re-Emerging Viral Infections. Front Immunol 9, 1379 (2018).

8. Y. Liu, D. Olagnier, R. Lin, Host and Viral Modulation of RIG-I-Mediated Antiviral Immunity. Front Immunol 7, 662 (2016).

9. M. L. Goulet, D. Olagnier, Z. Xu, S. Paz, S. M. Belgnaoui, E. I. Lafferty, V. Janelle, M. Arguello, M. Paquet, K. Ghneim, S. Richards, A. Smith, P. Wilkinson, M. Cameron, U. Kalinke, S. Qureshi, A. Lamarre, E. K. Haddad, R. P. Sekaly, S. Peri, S. Balachandran, R. Lin, J. Hiscott, Systems analysis of a RIG-I agonist inducing broad spectrum inhibition of virus infectivity. PLoS Pathog 9, e1003298 (2013).

10. M. Goldeck, M. Schlee, G. Hartmann, V. Hornung, Enzymatic synthesis and purification of a defined RIG-I ligand. Methods Mol Biol 1169, 15–25 (2014).

11. C. Grutzner, Y. P. Tan, P. Muller, T. M. Schlee-Guimaraes, M. Jentzsch, J. L. Schmid-Burgk, M. Renn, R. Behrendt, G. Hartmann, Synthetic RIG-I agonist-mediated cancer immunotherapy synergizes with MAP kinase inhibition against BRAF-mutated melanoma. Mol Ther Nucleic Acids 35, 102283 (2024).

12. S. Marx, B. M. Kummerer, C. Grutzner, H. Kato, M. Schlee, M. Renn, E. Bartok, G. Hartmann, RIG-I-induced innate antiviral immunity protects mice from lethal SARS-CoV-2 infection. Mol Ther Nucleic Acids 27, 1225–1234 (2022).

13. C. Coch, J. P. Stumpel, V. Lilien-Waldau, D. Wohlleber, B. M. Kummerer, I. Bekeredjian-Ding, G. Kochs, N. Garbi, S. Herberhold, C. Schuberth-Wagner, J. Ludwig, W. Barchet, M. Schlee, A. Hoerauf, F. Bootz, P. Staeheli, G. Hartmann, E. Hartmann, RIG-I Activation Protects and Rescues from Lethal Influenza Virus Infection and Bacterial Superinfection. Mol Ther 25, 2093–2103 (2017).

14. S. Lambing, Y. P. Tan, P. Vasileiadou, S. Holdenrieder, P. Muller, C. Hagen, S. Garbe, R. Behrendt, M. Schlee, J. G. van den Boorn, E. Bartok, M. Renn, G. Hartmann, RIG-I immunotherapy overcomes radioresistance in p53-positive malignant melanoma. J Mol Cell Biol 15, (2023).

15. L. S. U. Schwab, R. Farrukee, J. F. Eleouet, M. A. Rameix-Welti, S. L. Londrigan, A. G. Brooks, A. C. Hurt, C. Coch, T. Zillinger, G. Hartmann, P. C. Reading, Retinoic Acid-Inducible Gene I Activation Inhibits Human Respiratory Syncytial Virus Replication in Mammalian Cells and in Mouse and Ferret Models of Infection. J Infect Dis 226, 2079–2088 (2022).

16. L. S. U. Schwab, S. L. Londrigan, A. G. Brooks, A. C. Hurt, A. Sahu, Y. M. Deng, J. Moselen, C. Coch, T. Zillinger, G. Hartmann, P. C. Reading, Induction of Interferon-Stimulated Genes Correlates with Reduced Growth of Influenza A Virus in Lungs after RIG-I Agonist Treatment of Ferrets. J Virol 96, e0055922 (2022).

17. T. Saito, D. M. Owen, F. Jiang, J. Marcotrigiano, M. Gale, Jr., Innate immunity induced by composition-dependent RIG-I recognition of hepatitis C virus RNA. Nature 454, 523–527 (2008).

18. G. Schnell, Y. M. Loo, J. Marcotrigiano, M. Gale, Jr., Uridine composition of the poly-U/UC tract of HCV RNA defines non-self recognition by RIG-I. PLoS Pathog 8, e1002839 (2012).

19. B. S. Ulloa, I. Barber-Axthelm, B. Berube, M. Duthie, S. Reed, R. Savan, M. Gale, Jr., Synthetic RIG-I-Agonist RNA Induces Death of Hepatocellular Carcinoma Cells. J Interferon Cytokine Res 45, 119–132 (2025).

20. L. Martinez-Gil, P. H. Goff, R. Hai, A. Garcia-Sastre, M. L. Shaw, P. Palese, A Sendai virus-derived RNA agonist of RIG-I as a virus vaccine adjuvant. J Virol 87, 1290–1300 (2013).

21. W. Wang, A. M. Pyle, The RIG-I receptor adopts two different conformations for distinguishing host from viral RNA ligands. Mol Cell 82, 4131–4144 e4136 (2022).

22. M. M. Linehan, T. H. Dickey, E. S. Molinari, M. E. Fitzgerald, O. Potapova, A. Iwasaki, A. M. Pyle, A minimal RNA ligand for potent RIG-I activation in living mice. Sci Adv 4, e1701854 (2018).

23. D. Thoresen, W. Wang, D. Galls, R. Guo, L. Xu, A. M. Pyle, The molecular mechanism of RIG-I activation and signaling. Immunol Rev 304, 154–168 (2021).

24. X. Ren, M. M. Linehan, A. Iwasaki, A. M. Pyle, RIG-I Selectively Discriminates against 5’-Monophosphate RNA. Cell Rep 26, 2019–2027 e2014 (2019).

25. V. T. Mihaylova, Y. Kong, O. Fedorova, L. Sharma, C. S. Dela Cruz, A. M. Pyle, A. Iwasaki, E. F. Foxman, Regional Differences in Airway Epithelial Cells Reveal Tradeoff between Defense against Oxidative Stress and Defense against Rhinovirus. Cell Rep 24, 3000–3007 e3003 (2018).

26. X. Jiang, V. Muthusamy, O. Fedorova, Y. Kong, D. J. Kim, M. Bosenberg, A. M. Pyle, A. Iwasaki, Intratumoral delivery of RIG-I agonist SLR14 induces robust antitumor responses. J Exp Med 216, 2854–2868 (2019).

27. D. T. Thoresen, D. Galls, B. Gotte, W. Wang, A. M. Pyle, A rapid RIG-I signaling relay mediates efficient antiviral response. Mol Cell 83, 90–104 e104 (2023).

28. T. Mao, B. Israelow, C. Lucas, C. B. F. Vogels, M. L. Gomez-Calvo, O. Fedorova, M. I. Breban, B. L. Menasche, H. Dong, M. Linehan, S.-C.-G. S. I. Yale, C. B. Wilen, M. L. Landry, N. D. Grubaugh, A. M. Pyle, A. Iwasaki, A stem-loop RNA RIG-I agonist protects against acute and chronic SARS-CoV-2 infection in mice. J Exp Med 219, (2022).

29. R. Channappanavar, A. R. Fehr, R. Vijay, M. Mack, J. Zhao, D. K. Meyerholz, S. Perlman, Dysregulated Type I Interferon and Inflammatory Monocyte-Macrophage Responses Cause Lethal Pneumonia in SARS-CoV-Infected Mice. Cell Host Microbe 19, 181–193 (2016).

30. I. E. Galani, N. Rovina, V. Lampropoulou, V. Triantafyllia, M. Manioudaki, E. Pavlos, E. Koukaki, P. C. Fragkou, V. Panou, V. Rapti, O. Koltsida, A. Mentis, N. Koulouris, S. Tsiodras, A. Koutsoukou, E. Andreakos, Untuned antiviral immunity in COVID-19 revealed by temporal type I/III interferon patterns and flu comparison. Nat Immunol 22, 32–40 (2021).

31. S. Nair, Y. Wu, T. M. Nguyen, K. Fink, D. Luo, C. Ruedl, Intranasal Delivery of RIG-I Agonist Drives Pulmonary Myeloid Cell Activation in Mice. Front Immunol 13, 910192 (2022).

32. D. Olagnier, F. E. Scholte, C. Chiang, I. C. Albulescu, C. Nichols, Z. He, R. Lin, E. J. Snijder, M. J. van Hemert, J. Hiscott, Inhibition of dengue and chikungunya virus infections by RIG-I-mediated type I interferon-independent stimulation of the innate antiviral response. J Virol 88, 4180–4194 (2014).

33. H. Feldmann, A. Sprecher, T. W. Geisbert, Ebola. N Engl J Med 382, 1832–1842 (2020).

34. A. Markham, REGN-EB3: First Approval. Drugs 81, 175–178 (2021).

35. S. Mulangu, L. E. Dodd, R. T. Davey, Jr., O. Tshiani Mbaya, M. Proschan, D. Mukadi, M. Lusakibanza Manzo, D. Nzolo, A. Tshomba Oloma, A. Ibanda, R. Ali, S. Coulibaly, A. C. Levine, R. Grais, J. Diaz, H. C. Lane, J. J. Muyembe-Tamfum, P. W. Group, B. Sivahera, M. Camara, R. Kojan, R. Walker, B. Dighero-Kemp, H. Cao, P. Mukumbayi, P. Mbala-Kingebeni, S. Ahuka, S. Albert, T. Bonnett, I. Crozier, M. Duvenhage, C. Proffitt, M. Teitelbaum, T. Moench, J. Aboulhab, K. Barrett, K. Cahill, K. Cone, R. Eckes, L. Hensley, B. Herpin, E. Higgs, J. Ledgerwood, J. Pierson, M. Smolskis, Y. Sow, J. Tierney, S. Sivapalasingam, W. Holman, N. Gettinger, D. Vallee, J. Nordwall, P. C. S. Team, A Randomized, Controlled Trial of Ebola Virus Disease Therapeutics. N Engl J Med 381, 2293–2303 (2019).

36. E. Callaway, ’Make Ebola a thing of the past’: first vaccine against deadly virus approved. Nature 575, 425–426 (2019).

37. I. Messaoudi, G. K. Amarasinghe, C. F. Basler, Filovirus pathogenesis and immune evasion: insights from Ebola virus and Marburg virus. Nat Rev Microbiol 13, 663–676 (2015).

38. X. Liu, E. Speranza, C. Munoz-Fontela, S. Haldenby, N. Y. Rickett, I. Garcia-Dorival, Y. Fang, Y. Hall, E. G. Zekeng, A. Ludtke, D. Xia, R. Kerber, R. Krumkamp, S. Duraffour, D. Sissoko, J. Kenny, N. Rockliffe, E. D. Williamson, T. R. Laws, M. N’Faly, D. A. Matthews, S. Gunther, A. R. Cossins, A. Sprecher, J. H. Connor, M. W. Carroll, J. A. Hiscox, Transcriptomic signatures differentiate survival from fatal outcomes in humans infected with Ebola virus. Genome Biol 18, 4 (2017).

39. S. Reynard, A. Journeaux, E. Gloaguen, J. Schaeffer, H. Varet, N. Pietrosemoli, M. Mateo, N. Baillet, C. Laouenan, H. Raoul, J. Mullaert, S. Baize, Immune parameters and outcomes during Ebola virus disease. JCI Insight 4, (2019).

40. H. Li, J. V. Kim, B. S. Pickering, Henipavirus zoonosis: outbreaks, animal hosts and potential new emergence. Front Microbiol 14, 1167085 (2023).

41. R. Pelissier, M. Iampietro, B. Horvat, Recent advances in the understanding of Nipah virus immunopathogenesis and anti-viral approaches. F1000Res 8, (2019).

42. B. Kaza, H. C. Aguilar, Pathogenicity and virulence of henipaviruses. Virulence 14, 2273684 (2023).

43. C. Mathieu, V. Guillaume, A. Sabine, K. C. Ong, K. T. Wong, C. Legras-Lachuer, B. Horvat, Lethal Nipah virus infection induces rapid overexpression of CXCL10. PLoS One 7, e32157 (2012).

44. P. Lawrence, B. Escudero-Perez, Henipavirus Immune Evasion and Pathogenesis Mechanisms: Lessons Learnt from Natural Infection and Animal Models. Viruses 14, (2022).

45. W. H. Organization, Pathogens prioritization: a scientific framework for epidemic and pandemic research preparedness. World Health Organization: Geneva, Switzerland, (2024).

46. D. Wright, J. Kortekaas, T. A. Bowden, G. M. Warimwe, Rift Valley fever: biology and epidemiology. J Gen Virol 100, 1187–1199 (2019).

47. M. K. Rostal, P. N. Thompson, A. Anyamba, B. Bett, C. Cetre-Sossah, V. Chevalier, M. Guarido, W. B. Karesh, A. Kemp, A. D. LaBeaud, A. Lubisi, L. Matthews, V. Msimang, M. K. Njenga, N. Ross, D. Tumusiime, W. C. Wilson, J. Weyer, J. T. Paweska, R. Swanepoel, Rift Valley fever epidemiology: shifting the paradigm and rethinking research priorities. Lancet Planet Health 9, 101299 (2025).

48. T. G. Wanninger, D. E. Millian, O. A. Saldarriaga, J. Maruyama, T. Saito, R. A. Reyna, S. Taniguchi, E. Arroyave, M. E. Connolly, H. L. Stevenson, S. Paessler, Macrophage infection, activation, and histopathological findings in ebolavirus infection. Front Cell Infect Microbiol 12, 1023557 (2022).

49. U. Stroher, E. West, H. Bugany, H. D. Klenk, H. J. Schnittler, H. Feldmann, Infection and activation of monocytes by Marburg and Ebola viruses. J Virol 75, 11025–11033 (2001).

50. M. Gupta, S. Mahanty, R. Ahmed, P. E. Rollin, Monocyte-derived human macrophages and peripheral blood mononuclear cells infected with ebola virus secrete MIP-1alpha and TNF-alpha and inhibit poly-IC-induced IFN-alpha in vitro. Virology 284, 20–25 (2001).

51. J. Olejnik, A. Forero, L. R. Deflube, A. J. Hume, W. A. Manhart, A. Nishida, A. Marzi, M. G. Katze, H. Ebihara, A. L. Rasmussen, E. Muhlberger, Ebolaviruses Associated with Differential Pathogenicity Induce Distinct Host Responses in Human Macrophages. J Virol 91, (2017).

52. R. Wu, F. Chen, N. Wang, D. Tang, R. Kang, ACOD1 in immunometabolism and disease. Cell Mol Immunol 17, 822–833 (2020).

53. J. S. Towner, J. Paragas, J. E. Dover, M. Gupta, C. S. Goldsmith, J. W. Huggins, S. T. Nichol, Generation of eGFP expressing recombinant Zaire ebolavirus for analysis of early pathogenesis events and high-throughput antiviral drug screening. Virology 332, 20–27 (2005).

54. J. Dyall, B. J. Hart, E. Postnikova, Y. Cong, H. Zhou, D. M. Gerhardt, D. Freeburger, J. Michelotti, A. N. Honko, L. E. DeWald, R. S. Bennett, G. G. Olinger, Jr., P. B. Jahrling, L. E. Hensley, Interferon-beta and Interferon-gamma Are Weak Inhibitors of Ebola Virus in Cell-Based Assays. J Infect Dis 215, 1416–1420 (2017).

55. T. Klouda, Y. Hao, H. Kim, J. Kim, J. Olejnik, A. J. Hume, S. Ayyappan, X. Hong, J. Melero-Martin, Y. Fang, Q. Wang, X. Zhou, E. Muhlberger, H. Jia, R. F. Padera, Jr., B. A. Raby, K. Yuan, Interferon-alpha or -beta facilitates SARS-CoV-2 pulmonary vascular infection by inducing ACE2. Angiogenesis, (2021).

56. F. Villinger, P. E. Rollin, S. S. Brar, N. F. Chikkala, J. Winter, J. B. Sundstrom, S. R. Zaki, R. Swanepoel, A. A. Ansari, C. J. Peters, Markedly elevated levels of interferon (IFN)-gamma, IFN-alpha, interleukin (IL)-2, IL-10, and tumor necrosis factor-alpha associated with fatal Ebola virus infection. J Infect Dis 179 Suppl 1, S188–191 (1999).

57. J. Olejnik, A. J. Hume, D. W. Leung, G. K. Amarasinghe, C. F. Basler, E. Muhlberger, Filovirus Strategies to Escape Antiviral Responses. Curr Top Microbiol Immunol, (2017).

58. A. L. Hartman, L. Ling, S. T. Nichol, M. L. Hibberd, Whole-genome expression profiling reveals that inhibition of host innate immune response pathways by Ebola virus can be reversed by a single amino acid change in the VP35 protein. J Virol 82, 5348–5358 (2008).

59. W. A. Scoon, L. Mancio-Silva, E. L. Suder, C. Villacorta-Martin, J. Lindstrom-Vautrin, J. G. Bernbaum, S. Mazur, R. F. Johnson, J. Olejnik, E. Y. Flores, A. Mithal, F. Wang, A. J. Hume, J. E. Kaserman, S. March-Riera, A. A. Wilson, S. N. Bhatia, E. Muhlberger, G. Mostoslavsky, Ebola virus infection induces a delayed type I IFN response in bystander cells and the shutdown of key liver genes in human iPSC-derived hepatocytes. Stem Cell Reports 17, 2286–2302 (2022).

60. J. Elliff, H. Van Ert, K. Sevcik, M. A. Djurkovic, D. Rudd, F. Gourronc, A. Klingelhutz, O. Shtanko, W. Maury, Interferons Inhibit Ebola Virus Infection of Human Keratinocytes. Viruses 17, (2025).

61. E. Y. Flores, A. J. Hume, J. Olejnik, A. Mithal, A. D’Amico, M. Yang, P. Bawa, F. Wang, A. K. O’Connell, A. Tseng, N. A. Crossland, G. Mostoslavsky, E. Muhlberger, Filovirus infection disrupts epithelial barrier function and ion transport in human iPSC-derived gut organoids. PLoS Pathog 21, e1013698 (2025).

62. L. E. Hensley, H. A. Young, P. B. Jahrling, T. W. Geisbert, Proinflammatory response during Ebola virus infection of primate models: possible involvement of the tumor necrosis factor receptor superfamily. Immunol Lett 80, 169–179 (2002).

63. B. Escudero-Perez, V. A. Volchkova, O. Dolnik, P. Lawrence, V. E. Volchkov, Shed GP of Ebola virus triggers immune activation and increased vascular permeability. PLoS Pathog 10, e1004509 (2014).

64. M. Bray, T. W. Geisbert, Ebola virus: The role of macrophages and dendritic cells in the pathogenesis of Ebola hemorrhagic fever. Int J Biochem Cell Biol 37, 1560–1566 (2005).

65. A. J. Hume, B. Heiden, J. Olejnik, E. L. Suder, S. Ross, W. A. Scoon, E. Bullitt, M. Ericsson, M. R. White, J. Turcinovic, T. T. N. Thao, R. M. Hekman, J. E. Kaserman, J. Huang, K. D. Alysandratos, G. E. Toth, F. Jakab, D. N. Kotton, A. A. Wilson, A. Emili, V. Thiel, J. H. Connor, G. Kemenesi, D. Cifuentes, E. Muhlberger, Recombinant Lloviu virus as a tool to study viral replication and host responses. PLoS Pathog 18, e1010268 (2022).

66. A. Maisner, J. Neufeld, H. Weingartl, Organ- and endotheliotropism of Nipah virus infections in vivo and in vitro. Thromb Haemost 102, 1014–1023 (2009).

67. L. T. Ang, A. T. Nguyen, K. J. Liu, A. Chen, X. Xiong, M. Curtis, R. M. Martin, B. C. Raftry, C. Y. Ng, U. Vogel, A. Lander, B. J. Lesch, J. L. Fowler, A. R. Holman, T. Chai, S. Vijayakumar, F. P. Suchy, T. Nishimura, J. Bhadury, M. H. Porteus, H. Nakauchi, C. Cheung, S. C. George, K. Red-Horse, J. B. Prescott, K. M. Loh, Generating human artery and vein cells from pluripotent stem cells highlights the arterial tropism of Nipah and Hendra viruses. Cell 185, 2523–2541 e2530 (2022).

68. K. Meier, J. Olejnik, A. J. Hume, E. Muhlberger, A Comparative Assessment of the Pathogenic Potential of Newly Discovered Henipaviruses. Pathogens 13, (2024).

69. A. J. Hume, E. Muhlberger, Distinct Genome Replication and Transcription Strategies within the Growing Filovirus Family. J Mol Biol, (2019).

70. D. R. Smith, K. E. Steele, J. Shamblin, A. Honko, J. Johnson, C. Reed, M. Kennedy, J. L. Chapman, L. E. Hensley, The pathogenesis of Rift Valley fever virus in the mouse model. Virology 407, 256–267 (2010).

71. E. Haddock, H. Feldmann, A. Marzi, Ebola Virus Infection in Commonly Used Laboratory Mouse Strains. J Infect Dis 218, S453–S457 (2018).

72. S. S. Burkart, D. Schweinoch, J. Frankish, C. Sparn, S. Wust, C. Urban, M. Merlo, V. G. Magalhaes, A. Piras, A. Pichlmair, J. Willemsen, L. Kaderali, M. Binder, High-resolution kinetic characterization of the RIG-I-signaling pathway and the antiviral response. Life Sci Alliance 6, (2023).

73. W. B. Cardenas, Y. M. Loo, M. Gale, Jr., A. L. Hartman, C. R. Kimberlin, L. Martinez-Sobrido, E. O. Saphire, C. F. Basler, Ebola virus VP35 protein binds double-stranded RNA and inhibits alpha/beta interferon production induced by RIG-I signaling. J Virol 80, 5168–5178 (2006).

74. D. W. Leung, K. C. Prins, D. M. Borek, M. Farahbakhsh, J. M. Tufariello, P. Ramanan, J. C. Nix, L. A. Helgeson, Z. Otwinowski, R. B. Honzatko, C. F. Basler, G. K. Amarasinghe, Structural basis for dsRNA recognition and interferon antagonism by Ebola VP35. Nat Struct Mol Biol 17, 165–172 (2010).

75. P. Ramanan, M. R. Edwards, R. S. Shabman, D. W. Leung, A. C. Endlich-Frazier, D. M. Borek, Z. Otwinowski, G. Liu, J. Huh, C. F. Basler, G. K. Amarasinghe, Structural basis for Marburg virus VP35-mediated immune evasion mechanisms. Proc Natl Acad Sci U S A 109, 20661–20666 (2012).

76. S. Bale, J. P. Julien, Z. A. Bornholdt, C. R. Kimberlin, P. Halfmann, M. A. Zandonatti, J. Kunert, G. J. Kroon, Y. Kawaoka, I. J. MacRae, I. A. Wilson, E. O. Saphire, Marburg virus VP35 can both fully coat the backbone and cap the ends of dsRNA for interferon antagonism. PLoS Pathog 8, e1002916 (2012).

77. S. Bale, J. P. Julien, Z. A. Bornholdt, A. S. Krois, I. A. Wilson, E. O. Saphire, Ebolavirus VP35 coats the backbone of double-stranded RNA for interferon antagonism. J Virol 87, 10385–10388 (2013).

78. C. F. Basler, A. Mikulasova, L. Martinez-Sobrido, J. Paragas, E. Mühlberger, M. Bray, H. D. Klenk, P. Palese, A. Garcia-Sastre, The Ebola virus VP35 protein inhibits activation of interferon regulatory factor 3. J Virol 77, 7945–7956 (2003).

79. K. C. Prins, W. B. Cardenas, C. F. Basler, Ebola virus protein VP35 impairs the function of interferon regulatory factor-activating kinases IKKepsilon and TBK-1. J Virol 83, 3069–3077 (2009).

80. A. L. Hartman, B. H. Bird, J. S. Towner, Z. A. Antoniadou, S. R. Zaki, S. T. Nichol, Inhibition of IRF-3 activation by VP35 is critical for the high level of virulence of ebola virus. J Virol 82, 2699–2704 (2008).

81. K. C. Prins, S. Delpeut, D. W. Leung, O. Reynard, V. A. Volchkova, S. P. Reid, P. Ramanan, W. B. Cardenas, G. K. Amarasinghe, V. E. Volchkov, C. F. Basler, Mutations abrogating VP35 interaction with double-stranded RNA render Ebola virus avirulent in guinea pigs. J Virol 84, 3004–3015 (2010).

82. A. Pinski, C. Woolsey, A. Jankeel, R. Cross, C. F. Basler, T. Geisbert, I. Messaoudi, Transcriptional Analysis of Lymphoid Tissues from Infected Nonhuman Primates Reveals the Basis for Attenuation and Immunogenicity of an Ebola Virus Encoding a Mutant VP35 Protein. J Virol 95, (2021).

83. S. Baize, E. M. Leroy, A. J. Georges, M. C. Georges-Courbot, M. Capron, I. Bedjabaga, J. Lansoud-Soukate, E. Mavoungou, Inflammatory responses in Ebola virus-infected patients. Clin Exp Immunol 128, 163–168 (2002).

84. A. E. Tseng, M. Carossino, H. P. Gertje, A. K. O’Connell, S. Gummuluru, V. B. Kolachalama, U. B. R. Balasuriya, J. H. Connor, R. S. Bennett, D. X. Liu, L. E. Hensley, N. A. Crossland, Hepatic proinflammatory myeloid phenotypes are a hallmark of Ebola virus Kikwit pathogenesis in rhesus monkeys. Vet Pathol 60, 473–487 (2023).

85. S. Pattabhi, C. R. Wilkins, R. Dong, M. L. Knoll, J. Posakony, S. Kaiser, C. E. Mire, M. L. Wang, R. C. Ireton, T. W. Geisbert, K. M. Bedard, S. P. Iadonato, Y. M. Loo, M. Gale, Jr., Targeting Innate Immunity for Antiviral Therapy through Small Molecule Agonists of the RLR Pathway. J Virol 90, 2372–2387 (2016).

86. A. K. McElroy, J. R. Harmon, T. D. Flietstra, S. Campbell, A. K. Mehta, C. S. Kraft, M. G. Lyon, J. B. Varkey, B. S. Ribner, C. J. Kratochvil, P. C. Iwen, P. W. Smith, R. Ahmed, S. T. Nichol, C. F. Spiropoulou, Kinetic Analysis of Biomarkers in a Cohort of US Patients With Ebola Virus Disease. Clin Infect Dis 63, 460–467 (2016).

87. L. M. Smith, L. E. Hensley, T. W. Geisbert, J. Johnson, A. Stossel, A. Honko, J. Y. Yen, J. Geisbert, J. Paragas, E. Fritz, G. Olinger, H. A. Young, K. H. Rubins, C. L. Karp, Interferon-beta Therapy Prolongs Survival in Rhesus Macaque Models of Ebola and Marburg Hemorrhagic Fever. J Infect Dis, (2013).

88. C. F. Spiropoulou, P. Ranjan, M. B. Pearce, T. K. Sealy, C. G. Albarino, S. Gangappa, T. Fujita, P. E. Rollin, S. T. Nichol, T. G. Ksiazek, S. Sambhara, RIG-I activation inhibits ebolavirus replication. Virology 392, 11–15 (2009).

89. L. D. Jasenosky, C. Cadena, C. E. Mire, V. Borisevich, V. Haridas, S. Ranjbar, A. Nambu, S. Bavari, V. Soloveva, S. Sadukhan, G. H. Cassell, T. W. Geisbert, S. Hur, A. E. Goldfeld, The FDA-Approved Oral Drug Nitazoxanide Amplifies Host Antiviral Responses and Inhibits Ebola Virus. iScience 19, 1279–1290 (2019).

90. M. S. Park, M. L. Shaw, J. Munoz-Jordan, J. F. Cros, T. Nakaya, N. Bouvier, P. Palese, A. Garcia-Sastre, C. F. Basler, Newcastle disease virus (NDV)-based assay demonstrates interferon-antagonist activity for the NDV V protein and the Nipah virus V, W, and C proteins. J Virol 77, 1501–1511 (2003).

91. C. F. Basler, Nipah and hendra virus interactions with the innate immune system. Curr Top Microbiol Immunol 359, 123–152 (2012).

92. N. Becker, A. Maisner, Nipah Virus Impairs Autocrine IFN Signaling by Sequestering STAT1 and STAT2 into Inclusion Bodies. Viruses 15, (2023).

93. L. G. Thorne, A. K. Reuschl, L. Zuliani-Alvarez, M. V. X. Whelan, J. Turner, M. Noursadeghi, C. Jolly, G. J. Towers, SARS-CoV-2 sensing by RIG-I and MDA5 links epithelial infection to macrophage inflammation. EMBO J 40, e107826 (2021).

94. A. Billecocq, M. Spiegel, P. Vialat, A. Kohl, F. Weber, M. Bouloy, O. Haller, NSs protein of Rift Valley fever virus blocks interferon production by inhibiting host gene transcription. J Virol 78, 9798–9806 (2004).

95. C. Lei, J. Yang, J. Hu, X. Sun, On the Calculation of TCID(50) for Quantitation of Virus Infectivity. Virol Sin 36, 141–144 (2021).

96. B. H. Bird, C. G. Albarino, S. T. Nichol, Rift Valley fever virus lacking NSm proteins retains high virulence in vivo and may provide a model of human delayed onset neurologic disease. Virology 362, 10–15 (2007).

97. F. Wincott, A. DiRenzo, C. Shaffer, S. Grimm, D. Tracz, C. Workman, D. Sweedler, C. Gonzalez, S. Scaringe, N. Usman, Synthesis, deprotection, analysis and purification of RNA and ribozymes. Nucleic Acids Res 23, 2677–2684 (1995).

98. S. Andrews. (2017).

99. A. Dobin, C. A. Davis, F. Schlesinger, J. Drenkow, C. Zaleski, S. Jha, P. Batut, M. Chaisson, T. R. Gingeras, STAR: ultrafast universal RNA-seq aligner. Bioinformatics 29, 15–21 (2013).

100. Y. Liao, G. K. Smyth, W. Shi, featureCounts: an efficient general purpose program for assigning sequence reads to genomic features. Bioinformatics 30, 923–930 (2014).

101. M. D. Robinson, D. J. McCarthy, G. K. Smyth, edgeR: a Bioconductor package for differential expression analysis of digital gene expression data. Bioinformatics 26, 139–140 (2010).

102. M. E. Ritchie, B. Phipson, D. Wu, Y. Hu, C. W. Law, W. Shi, G .K. Smyth, limma powers differential expression analyses for RNA-sequencing and microarray studies. Nucleic Acids Res 43, e47 (2015).

103. S. Su, C. W. Law, C. Ah-Cann, M. L. Asselin-Labat, M. E. Blewitt, M. E. Ritchie, Glimma: interactive graphics for gene expression analysis. Bioinformatics 33, 2050–2052 (2017).

104. A. Sergushichev, An algorithm for fast preranked gene set enrichment analysis using cumulative statistic calculation. BioRxiv 60012, 1–9 (2016).

105. P. Bankhead, M. B. Loughrey, J. A. Fernandez, Y. Dombrowski, D. G. McArt, P. D. Dunne, S. McQuaid, R. T. Gray, L. J. Murray, H. G. Coleman, J. A. James, M. Salto-Tellez, P. W. Hamilton, QuPath: Open source software for digital pathology image analysis. Sci Rep 7, 16878 (2017).

106. B. B. Gowen, M. H. Wong, K. H. Jung, A. B. Sanders, M. Mendenhall, K. W. Bailey, Y. Furuta, R. W. Sidwell, In vitro and in vivo activities of T-705 against arenavirus and bunyavirus infections. Antimicrob Agents Chemother 51, 3168–3176 (2007).

107. L. J. Reed, H. Muench, A Simple Method of Estimating Fifty Per Cent Endpoints12. American Journal of Epidemiology 27, 493–497 (1938).

108. A. Marzi, S. Chadinah, E. Haddock, F. Feldmann, N. Arndt, C. Martellaro, D. P. Scott, P. W. Hanley, T. G. Nyenswah, S. Sow, M. Massaquoi, H. Feldmann, Recently Identified Mutations in the Ebola Virus-Makona Genome Do Not Alter Pathogenicity in Animal Models. Cell Rep 23, 1806–1816 (2018).

